# Interplay of p53 and XIAP protein dynamics orchestrates cell fate in response to chemotherapy

**DOI:** 10.1101/2022.12.07.519451

**Authors:** Roba Abukwaik, Elias Vera-Siguenza, Daniel A. Tennant, Fabian Spill

**Author notes:** Correspondence should be addressed to Fabian Spill;, Elias Vera-Siguenza;, Roba Abukwaik.

## Abstract

Chemotherapeutic drugs are used to treat almost all types of cancer, but the intended response, i.e., elimination, is often incomplete, with a subset of cancer cells resisting treatment. Two critical factors play a role in chemoresistance: the p53 tumour suppressor gene and the X-linked inhibitor of apoptosis (XIAP). These proteins have been shown to act synergistically to elicit cellular responses upon DNA damage induced by chemotherapy, yet, the mechanism is poorly understood. This study introduces a mathematical model characterising the apoptosis pathway activation by p53 before and after mitochondrial outer membrane permeabilisation upon treatment with the chemotherapy Doxorubicin (Dox). *“In-silico”* simulations show that the p53 dynamics change dose-dependently. Under medium to high doses of Dox, p53 concentration ultimately stabilises to a high level regardless of XIAP concentrations. However, caspase-3 activation may be triggered or not depending on the XIAP induction rate, ultimately determining whether the cell will perish or resist. Consequently, the model predicts that failure to activate apoptosis in some cancer cells expressing wild-type p53 might be due to heterogeneity between cells in upregulating the XIAP protein, rather than due to the p53 protein concentration. Our model suggests that the interplay of the p53 dynamics and the XIAP induction rate is critical to determine the cancer cells’ therapeutic response.

## 1. Introduction

The last few decades have seen an increase in novel cancer treatments, including immuno- and targeted therapies. Despite these advances, traditional chemotherapy is still one of the prevailing methods to manage this incurable group of diseases [1]. The desired goal of chemotherapy is to eliminate all cancer cells, yet the malignancy’s ability to escape apoptosis (a major form of programmed cell death) plays a crucial role in promoting resistance to chemotherapy, limiting its efficacy [1, 2]. Increasing the dose may help eliminate more cancer cells. However, doses are limited by toxicity to other tissues as they preferentially, but not exclusively, target the rapidly dividing cancer cells [1, 2].

The standard class of chemotherapy drugs called “genotoxic agents”, examples of which include Cisplatin, Doxorubicin and Etoposide, depend on the p53 protein to trigger apoptosis by inducing DNA damage [2, 3, 4, 5, 6]. Cells with DNA damage activate kinases that disrupt the interaction between p53 and its negative regulator murine double minute 2 (Mdm2), leading to stabilisation and accumulation of p53 protein [3, 7, 8, 9]. Consequently, p53 commences transcriptional activation of genes involved in cell cycle arrest and apoptosis, determining the cell’s fate [9].

Unfortunately, clinical evidence has shown that patients undergoing genotoxic chemotherapy display variable degrees of chemoresistance, contributing to relapse or metastasis [1, 2, 3, 5]. Thus, understanding how p53 orchestrates cell fate decisions is crucial to improve patient prognosis.

Previous studies have assumed that p53-mediated cell fate decisions depend on the higher affinity of p53 for pro-arrest target gene promoters compared to pro-apoptotic genes; the so-called “affinity model”. This model suggests that low p53 levels (“level” denoting concentration) preferentially transactivate pro-arrest target genes, whereas high p53 levels are necessary to transactivate pro-apoptotic targets [10, 11]. However, this idea has been contested as many subsequent studies asserted that protein levels of p53 targets increase proportionally to p53 expression, irrespective of their function in arrest or apoptosis [6, 12]. Kracikova et al. examined the hypothesis by testing selected pro-arrest and pro-apoptotic p53 target genes induced with low and high p53 concentrations in non-tumorigenic wild-type p53 human mammary epithelial cells. Their results showed that cells contained both pro-arrest and -apoptotic genes with no evidence of differential upregulation by low and high p53 levels [12]. In agreement, Chen et al. compared the expression levels of well-characterised p53 target genes involved in cell-cycle arrest and apoptosis from cells treated with low and high chemotherapy doses of Etoposide [6]. Their experiments showed that both pro-arrest and -apoptotic genes were significantly induced under low and high doses, indicating no differential transactivation of pro-arrest and -apoptotic promoters. Alternatively, it has been proposed that p53 expression levels and its targets must overtake a so-called “apoptotic threshold” for a sustained period to trigger apoptosis. Anything below this threshold could be sufficient to induce arrest but not apoptosis [12].

Other studies have confirmed that the choice between cell-cycle arrest and apoptosis does not exclusively depend on p53 protein levels but also its dynamics. This is because different stresses elicit different temporal profiles of p53, indicating that temporal p53 dynamics play a role in the response mechanism [13, 14]. This hypothesis has been corroborated by experiments demonstrating how p53 oscillations lead to DNA repair and cell-cycle arrest, whereas cells with high stable p53 levels undergo senescence and apoptosis [13, 14]. It has also been shown that p53 dynamics can switch between modes in a damage/dose-dependent manner. Exposure to low-dose chemotherapy triggers a p53 pulsatile (oscillatory) behaviour which signals the cell cycle arrest fate. Conversely, high doses of chemo-drugs induce sustained activation of p53, provoking apoptosis [4, 6]. These observations point to a mechanism whereby p53 oscillations (under minor DNA damage) maintain p53 and its target genes at a low level, sufficient for cell cycle progression inhibition. In contrast, the monotonic p53 (under acute DNA damage) instigates robust induction of pro-apoptotic genes to execute apoptosis.

Recent experiments by Paek et al.,(2016) found that maximum p53 levels did not vary between apoptotic and surviving cells when quantifying p53 levels in human colon cancer cells treated with Cisplatin. Their results indicate that cell death is not simply determined by a fixed p53 threshold [5]. Instead, the upregulation of X-linked inhibitor of apoptosis protein (XIAP) and other apoptotic inhibitor proteins in response to chemotherapy could prevent caspases from activating apoptosis, despite high levels of p53. The observations were further corroborated by Holcik et al., who demonstrated that ionising irradiation induces upregulation of XIAP in human lung cancer cells that correlates with increased resistance to radiation [15]. Many other studies confirm alterations of XIAP in various human cancers with poor prognosis and resistance [16, 17].

Although many individual molecules involved in cell fate decisions have been identified, the interactions of these molecules on different timescales remain largely unexplored. In this study, we offer to leverage the predictive power of mathematics to underpin and elucidate those difficult-to-see mechanisms behind the machinery driving cell fate. To do so, we constructed a mathematical model of the apoptosis pathway activation by p53 under Doxorubicin (Dox) treatment. Our study is based on previous experimental and modelling efforts, and it expands those to analyse and predict the impact XIAP up-regulation has following chemotherapy on the fractional killing of cancer cells. In general, this study is not focused on a specific type of cancer but may mimic the type of cancers that see upregulation in the XIAP protein following chemotherapy drugs. Given that mitochondrial outer membrane permeabilisation (MOMP) is typically considered the “point of no return” in the apoptosis pathway, our model results argue that cell response heterogeneity could be linked to apoptosis impairment post-MOMP event [5, 16, 17, 18, 19].

## 2. Model and Assumptions

Our model comprises two compartments and ten key molecular species in the apoptosis pathway with positive and negative feedback loops schematically shown in Fig. 1. The compartments of our model are the cell’s cytoplasm and its nucleus. The cytoplasmic and nuclear molecular species are represented by the subscripts ‘c’ and ‘n’, respectively. Additionally, the ‘*’ superscript symbol is used to denote active species in the case of species that exist in two states (active and inactive). The presence of some molecules in one of these compartments mainly affects their activities and the downstream signals in the apoptosis pathway. The following section briefly discusses the cellular events within these compartments in response to chemotherapy Dox.

**Figure 1:**
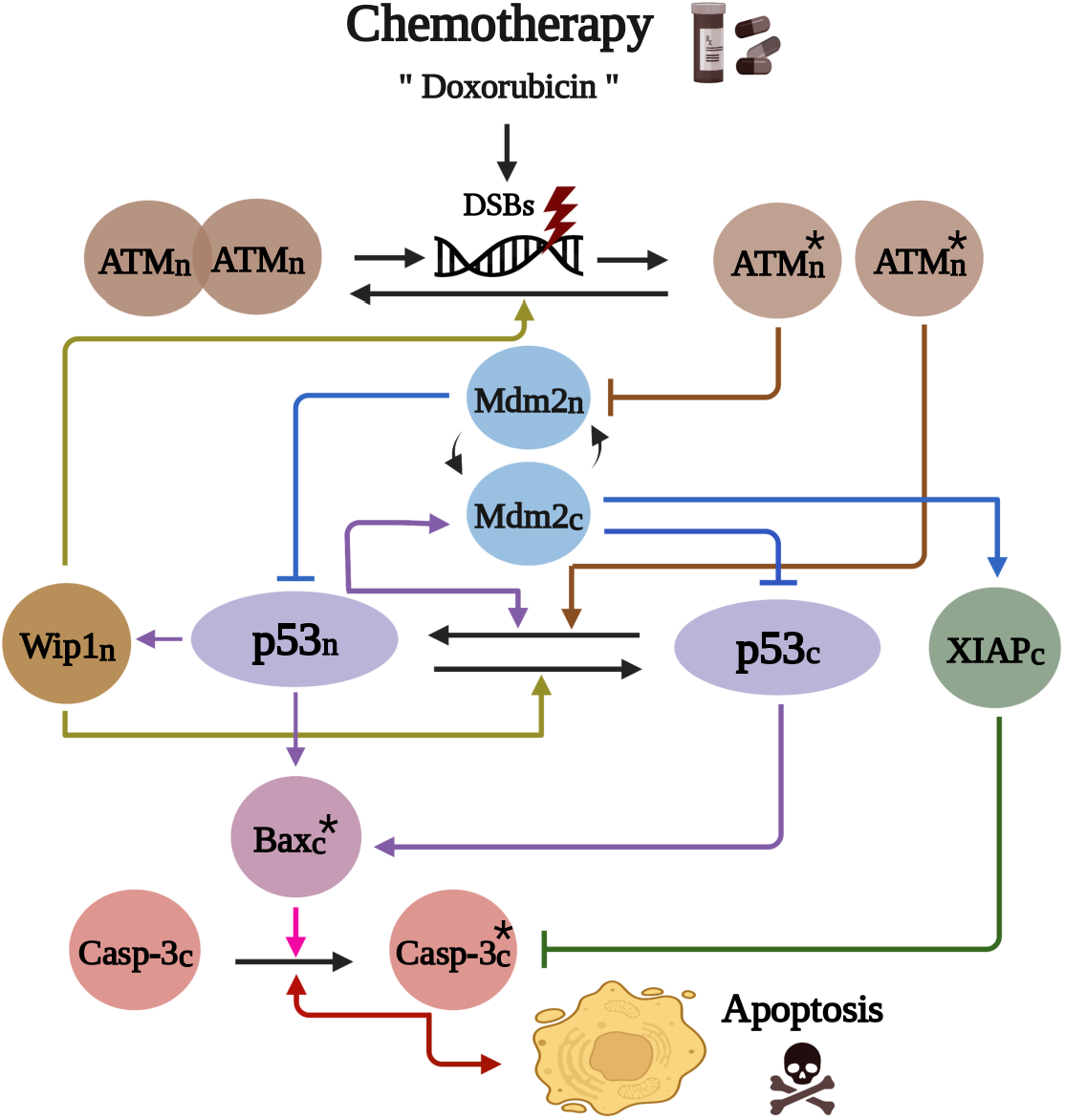
Schematic diagram depicting the signalling pathway of key molecules involved in the apoptosis pathway in response to DNA double-strand breaks (DSBs) following chemotherapy (Doxorubicin). Here we denote active nuclear ATM 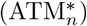, cytoplasmic p53 (p53_*c*_), nuclear p53 (p53_*n*_), cytoplasmic Mdm2 (Mdm2_*c*_), nuclear Mdm2 (Mdm2_*n*_), nuclear Wip1 (Wip1_*n*_), active cytoplasmic Bax 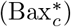, cytoplasmic XIAP (XIAP_*c*_), inactive cytoplasmic caspase-3 (Casp-3_*c*_), and active cytoplasmic caspase-3 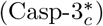.

### 2.1. Model Assumptions

#### 2.1.1. p53 and Mdm2 interaction

In normal cells, p53 protein is produced constantly from tumour suppressor gene TP53 [20]. Nevertheless, p53 protein levels are kept low by the negative regulator, Mdm2. Mdm2 is an E3 ubiquitin ligase that regulates the p53 degradation via the ubiquitin-proteasome pathway [21, 22]. However, Mdm2 production is also controlled by p53. If the p53 level rises high, it induces the production of more Mdm2, forming a negative feedback loop that drives the p53 levels back down [21, 22, 23].

#### 2.1.2. ATM activation

The occurrence of DNA double-strand breaks (DSBs) by genotoxic agents in chemotherapy initiates the activation of the protein kinase ataxia-telangiectasia mutated (ATM) [4, 24, 25]. ATM exists as a dimer in unperturbed cells. However, in response to DSBs, ATM is disassociated into active monomers through intermolecular phosphorylation [26]. Soon after, ATM phosphorylates p53 and Mdm2, influencing their stability [27]. ATM is reported to localise predominantly within the nucleus, with only small amounts residing in the cytoplasm [28]. However, the fraction of cytoplasmic ATM molecules has no kinase activity on p53 nor Mdm2 following DNA damage [28]. Accordingly, we assume that ATM and its active form are strictly nuclear proteins. Hence, p53 and Mdm2 phosphorylation by ATM is also rigorously nuclear.

#### 2.1.3. ATM activation affects p53 and Mdm2 and their location

p53 primarily accumulates in the cytoplasm in the basal state, where they interact with Mdm2 restricting them to the cytoplasm [29, 30]. Upon DNA damage, active ATM activates p53, leading to p53 accumulation and stabilisation via phosphorylation and acetylation. Increasing p53 acetylation levels promotes p53 nuclear translocation, where it can function as a transcription factor [4, 27]. Concomitantly, p53 phosphorylation on serine 15, or 20 supports p53 accumlation by disrupting binding to its regulator Mdm2 [7, 8, 27]. Furthermore, it has been shown that under normal conditions, p53 is present mainly as dimers (59%), followed by monomers (28%), and lastly, tetramers (13%) [31]. However, in response to DNA damage, ATM kinases can also mediate p53 phosphorylation on serine 392, promoting the tetramer formation of p53, which is essential for the p53 ability to transcribe genes [31, 32, 33]. Consequently, in our model, p53 is considered a cytoplasmic protein under unstressed conditions, but its active form is translocated into the nucleus by ATM activation, where it preferentially forms tetramers, binds DNA and transcriptionally acts as a tetramer.

It has been demonstrated that p53 is likewise subject to nuclear export. The nuclear export signal (NES) within the p53 protein is assumed to be necessary and sufficient to direct p53 nuclear export. However, the first p53’s NES is contained within the tetramerisation domain, blocked as long as p53 forms a tetramer. The second NES is located in the Mdm2-binding domain that can be attenuated via phosphorylation at serine 15 or 20 by ATM in the nucleus [29, 34]. Thus, phosphorylation and tetramerisation can inhibit p53 nuclear export by masking the NESs.

When it comes to Mdm2, it is preferentially located in the cytoplasm of unstressed cells [34], yet, the survival signalling (PI3K/Akt) promotes its translocation from the cytoplasm into the nucleus, inhibiting the p53 function [35, 36]. Accordingly, Mdm2 is assumed to be mainly cytoplasmic in our model, but might migrate to the nucleus due to survival signalling. Furthermore, upon DNA damage, active ATM induces phosphorylation of Mdm2, resulting in decreased activity and stability of Mdm2 protein [27].

#### 2.1.4. Activating p53 target genes

Due to p53 activation, p53 protein will commence gradual accumulation. As nuclear p53 levels increase, activation of all its target genes co-occurs, regardless of their functions on cell cycle arrest or apoptosis [12]. When this ensues, relatively low levels of p53 pro-arrest genes are sufficient to trigger cell growth arrest. But to enact apoptosis, the cell requires high concentrations of pro-apoptotic protein “Bax” to outweigh the anti-apoptotic protein “Bcl-2” and then start creating holes in the mitochondrial outer membrane [6, 18].

#### 2.1.5. Activating genes that inhibit p53 activation

Some p53 target genes can inhibit the p53 activation, such as Mdm2 and wild-type p53-induced phosphatase 1 (Wip1) [13]. High levels of Mdm2 increases the interaction rate between p53 and Mdm2, reducing the p53 stability [21, 22]. In addition, Mdm2 can affect p53 nucleocytoplasmic transportation. It has been reported that the Mdm2/p53 interaction in the nucleus shuttles p53 to the cytoplasm, where Mdm2 mediates the p53 degradation [29]. However, for simplicity, our model assumes that any interaction between Mdm2 and p53 will cause p53 degradation regardless of its location.

Wip1 has been found to reside in the nucleus exclusively [37], where it dephosphorylates ATM and p53, making p53 more vulnerable to Mdm2 and disabling ATM from phosphorylating p53 [13]. On the other hand, as mentioned above, phosphorylation of p53, whether on serine 15, 20 or 392, helps block its NESs from binding their receptors [29, 34]. Thus, the dephosphorylation of p53 by Wip1 can turn on p53 nuclear export by unmasking the NESs. Therefore, in our model, dephosphorylated p53 in the nucleus is assumed to be repositioned to the cytoplasm. These dynamics form negative feedback loops that prevent p53 levels from getting high enough to induce apoptosis [13].

#### 2.1.6. Activating genes that promote p53 activation

p53 also induces genes that promote its activation. Molecules such as the phosphatase with tensin homology (PTEN) inhibit translocation of Mdm2 into the nucleus and sustain p53 function. The protection of p53 from Mdm2 by PTEN forms a positive feedback loop that supports p53 activity and p53 accumulation [35, 38]. We include this positive feedback in our model as p53 activates itself.

#### 2.1.7. Caspase-3 and XIAP interaction following MOMP

Suppose the p53 level goes high enough to induce enough pro-apoptotic protein “Bax”. In that case, MOMP will occur, causing the release of the key mediators of intrinsic apoptosis, cytochrome c and the second mitochondria-derived activator of caspases (Smac), from the mitochondria to the cytoplasm. Following that, cytochrome c will start recruiting caspase-9 leading to caspase-3 activation [39]. However, starting activating caspase-3 or -9 does not mean the cell is undergoing apoptosis yet, as they will be targeted by the inhibitors of apoptosis proteins (IAPs) that inhibit caspase activation [16, 17, 40].

Some IAPs can bind and promote the ubiquitination of apoptotic caspases in vitro. However, out of the eight IAPs, XIAP is considered the main direct inhibitor of caspase activity and is the only member of this family able to directly inhibit both the initiation and execution phase of the caspase cascade [16, 41, 42]. Accordingly, in our model, we only consider the XIAP as a caspases inhibitor for simplicity.

The expression level of XIAP has been shown to be regulated at the translational level under apoptotic conditions, allowing for a rapid response to conditions in which XIAP function is required [43, 44]. The upregulation of XIAP protein levels is controlled by Mdm2 which was found to interact physically with the mRNA of XIAP and positively regulate XIAP IRES activity, increasing XIAP translation levels [45]. Therefore, we assume that Mdm2 would induce the XIAP protein in the cytoplasm under apoptotic conditions.

After MOMP, Smac is released from the mitochondria to support caspase activation by competing with caspases for XIAP interaction [46]. However, the upregulation of XIAP by Mdm2 increases the pool of XIAP available that might ultimately block caspase activation, disrupting the apoptotic program [5, 43]. Lastly, for the sake of simplicity, some intermediate effectors between the MOMP event and caspase-3 activation, like cytochrome c, Smac, and caspase-9, are not explicitly involved in our model (Fig. 1).

### 2.2. Model Equations

Based on the above assumptions, our model network spans two compartments, cytoplasm and nucleus, where directional fluxes determine the rate of change in concentration, with respect to time, of ten signalling molecules in the apoptosis pathway (Fig. 1).

Our model is closed; no molecular species escapes to the interstitial space (outside the cell). Hence, all signalling dynamics are limited to the cytoplasm and nucleus. Although we know these assumptions to be an approximation, our model provides a necessary first step towards constructing a more complex model that considers an open-cell model that incorporates the complexities of ion transport through the plasma membrane and other sub-cellular pathway organisation.

The variables of the model are the active nuclear ATM kinase 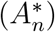, cytoplasmic p53 (*P_c_*), nuclear p53 (*P_n_*), cytoplasmic Mdm2 (*M_c_*), nuclear Mdm2 (*M_n_*), nuclear Wip1 (*W_n_*), cytoplasmic active Bax 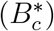, cytoplasmic XIAP (*X_c_*), inactive cytoplasmic caspase-3 (*C_3_c__*), active cytoplasmic caspase-3 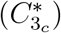, and chemotherapy Doxorubicin (*Dox*). Therefore, our system is comprised of eleven differential equations given by:

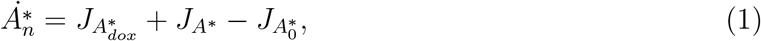

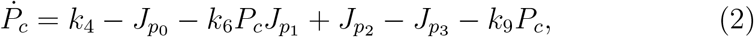

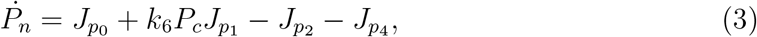

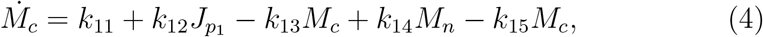

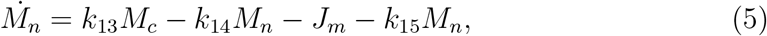

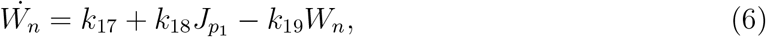

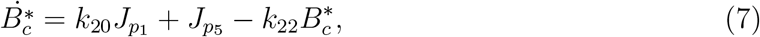

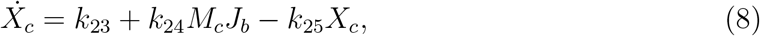

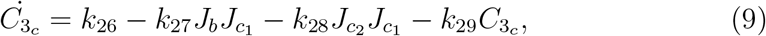

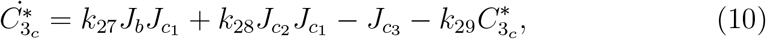

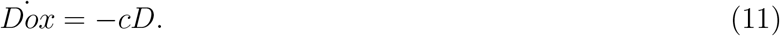

Each flux in the model, denoted by ‘J’, is represented by a mathematical sub-model based on experimental observations and previous mathematical models. These sub-models, and a detailed description of each, can be found in the Appendix section (Appendix A.1) of this article. A table of parameters, and initial conditions, can also be found in the Appendix section under Tables A.1 and A.2.

## 3. Results

Following chemotherapy treatments, cells trigger their self-defence machinery by stabilising and activating p53 to function primarily as a transcription factor, regulating the expression of downstream target genes that induce responses ranging from cell cycle arrest, DNA repair, and apoptosis. To scrutinise the dynamical responses of p53, we conducted a variety of “in-silico experimentation”, simulating cellular exposure to different doses of Dox. All our numerical simulations were carried out using the ode routine in MATLAB and Gear’s method in the XPPAUT software.

### 3.1. p53 exhibits various signalling modes under Doxorubicin treatment

After exposing the cell to low Dox levels, the protein concentration of active ATM, Mdm2, Wip1, and active Bax exhibits an oscillatory dynamics response that prompts a series of p53 pulses with fixed amplitude and period (Fig. 2a). The simulation result reflects how ATM is activated by sensing the DNA damage caused by chemotherapy. Soon after, phosphorylated p53 accumulates in the nucleus and transcriptionally activates its target genes. Then, Wip1 and Mdm2 protein levels increase to inhibit ATM and p53 activities, dropping their levels back again to basal levels. However, the residual DNA damage will reactivate ATM, producing another pulse of proteins and continuing the cycle (as above) until all damage is repaired. Notably, under this paradigm, cells undergo cell cycle arrest but not apoptosis as the caspase-3 activation levels remain low. Our result agrees with experimental findings that show pulsatile dynamics of p53 that exhibit constant mean amplitude and period in response to low Dox dosages, leading to cell cycle arrest [4].

**Figure 2:**
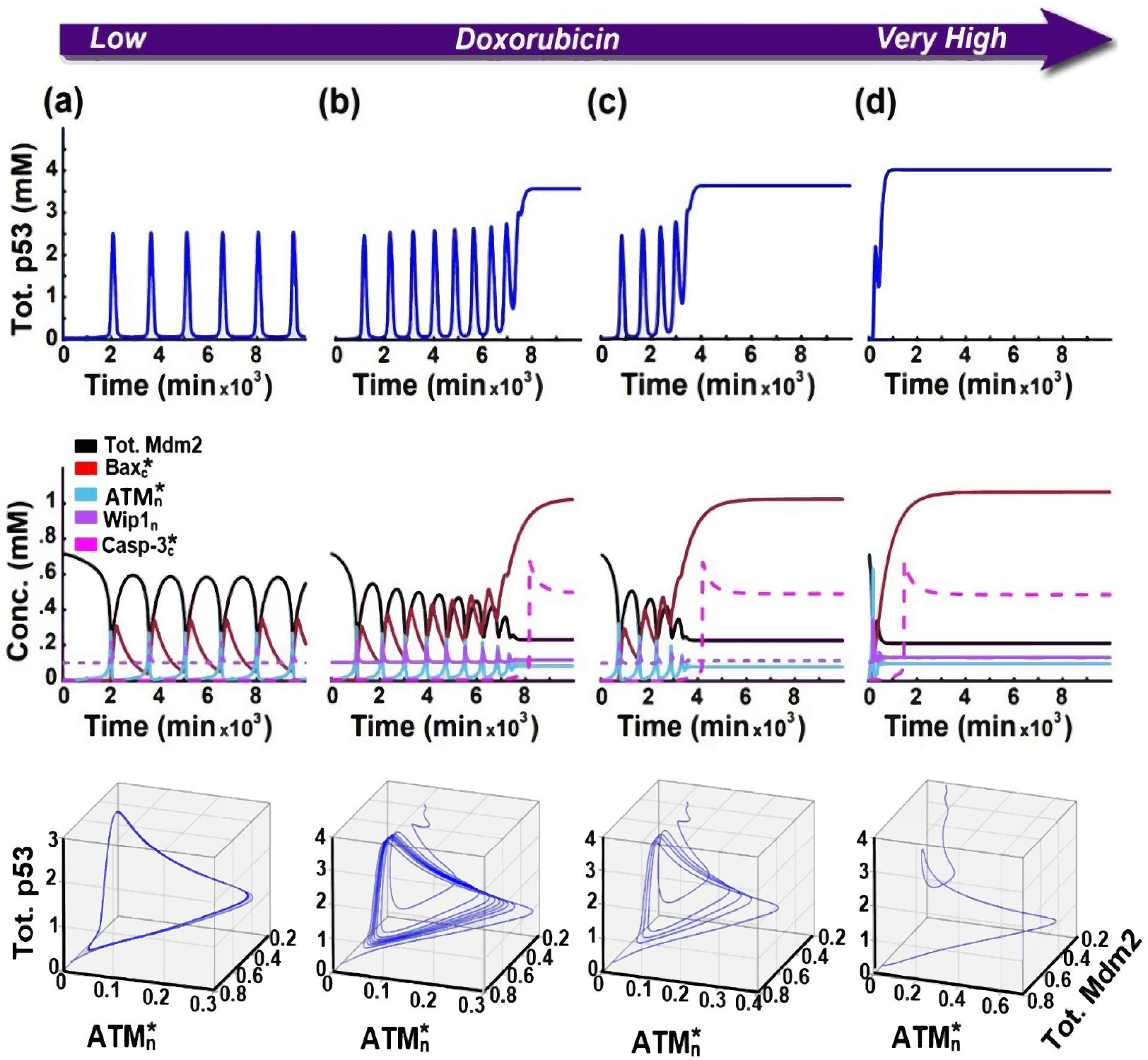
Model’s response to a low-to-very high Doxorubicin dosage. **(a)** (first column) Under low levels of Dox, cells converge to a stable limit cycle without triggering apoptosis. The system exhibits pulsatile behaviour by p53 to indicate cell repair, whereas the active caspase-3 concentrations remain low. **(b)** and **(c)** (second and third column) In response to medium Dox dosages, the solution’s limit cycle transitions from stable to unstable. This forces the system’s solutions toward the apoptosis steady state, reflected by a high concentration of active caspase-3. Under this paradigm, p53 displays biphasic dynamics; it initiates several oscillations, then jumps to approach a high-amplitude terminal pulse, executing apoptosis. **(d)** (last column) Upon relatively high doses of Dox, the concentration of p53 reaches a high concentration plateau, skipping any pulsatile behaviour. This behaviour prompts a fast convergence to the apoptosis steady state companioned by a fast activation of caspases-3.

In contrast, at moderate Dox doses, cells exhibit biphasic dynamics. p53 first initiates a series of oscillatory pulses but then jumps to a significantly high level, reaching a plateau called a terminal pulse. Here, the concentration of p53 is sufficiently elevated to trigger high enough levels of pro-apoptotic protein Bax followed by rapid caspase-3 activation to instigate apoptosis (Fig. 2b). In our model, the higher the Dox dose, the faster the transition to the second phase with the terminal pulse of p53, which is necessary to induce apoptosis (Figs. 2b, 2c). Our results are consistent with the experimental observations that show a duel-phase response of p53 at a moderate dose of Dox, where the appearance of p53 terminal pulse increases dose-dependently and is a direct indicator of whether or not apoptosis is executing [4].

On the other hand, cells with acute doses of Dox display a sustained induction of p53 without the first phase of discrete oscillations, activating caspase-3 much faster (Fig. 2d). Despite the delayed induction of pro-apoptotic protein Bax under moderate dosages, no significant difference was observed in its level induced by an acute dose of Dox, which is in further agreement with experimental data represented in [4].

When comparing all Dox dosage paradigms (low to high), we observe that cells change the p53 dynamics in a dose-dependent manner. Cells treated with low levels of Dox show single-phase pulsatile behaviours of p53 without the terminal pulse. Here, the minor damage will be sensed by ATM kinase and then convergence to a stable limit cycle without triggering the death fate (Fig. 2a). However, cells with moderate doses initiate sequential pulses of p53 followed by a high-amplitude terminal pulse, executing the cell death program. In this case, the limit cycle becomes unstable, pushing the system toward the apoptosis steady state with a high level of active caspase-3 (Figs. 2b, 2c). Under an acute dose of Dox, p53 displays a sustained activation, converging to apoptosis steady state faster and activating apoptosis more quickly (Fig. 2d).

### 3.2. Survival is accessible before apoptosis is initiated

Dox is totally removed or cleared from the cell by the time. Its elimination from tissues has a terminal half-life between 20 to 48 hours [47, 48]. Therefore, it is worth clarifying how the cell behaves when Dox gets cleared, whether before or after triggering apoptosis. To do so, we treat Dox concentration as a model variable (not as a parameter). Here, we assume that Dox is undergoing an exponential decay with a clearance rate *c*, and *D* is the initial dose level, which is considered a model input in our system (Eq. 11).

Our simulations revealed that cells under low doses of Dox (0.005 mM ≤ D < 0.065 mM) initiate a number of p53 pulses positively correlated with the dose level. Then, the p53 concentration is dropped back to its normal level, returning all other proteins to their basal concentrations, indicating a return to a survival steady state (Figs. 3b, 3c). On the contrary, when cells are challenged with moderate doses of Dox (0.065 mM ≤ D ≤ 1 mM), they initially vary the p53 level in a series of discrete pulses but then trigger the terminal pulse, directing the cell to the apoptosis steady state with a high level of active caspase-3 to guarantee apoptotic fate (Figs. 3d, 3e). However, after an acute application of Dox (*D* > 1 mM), the p53 level increases relatively faster, achieving the apoptosis steady state without oscillations (Fig. 3f).

**Figure 3:**
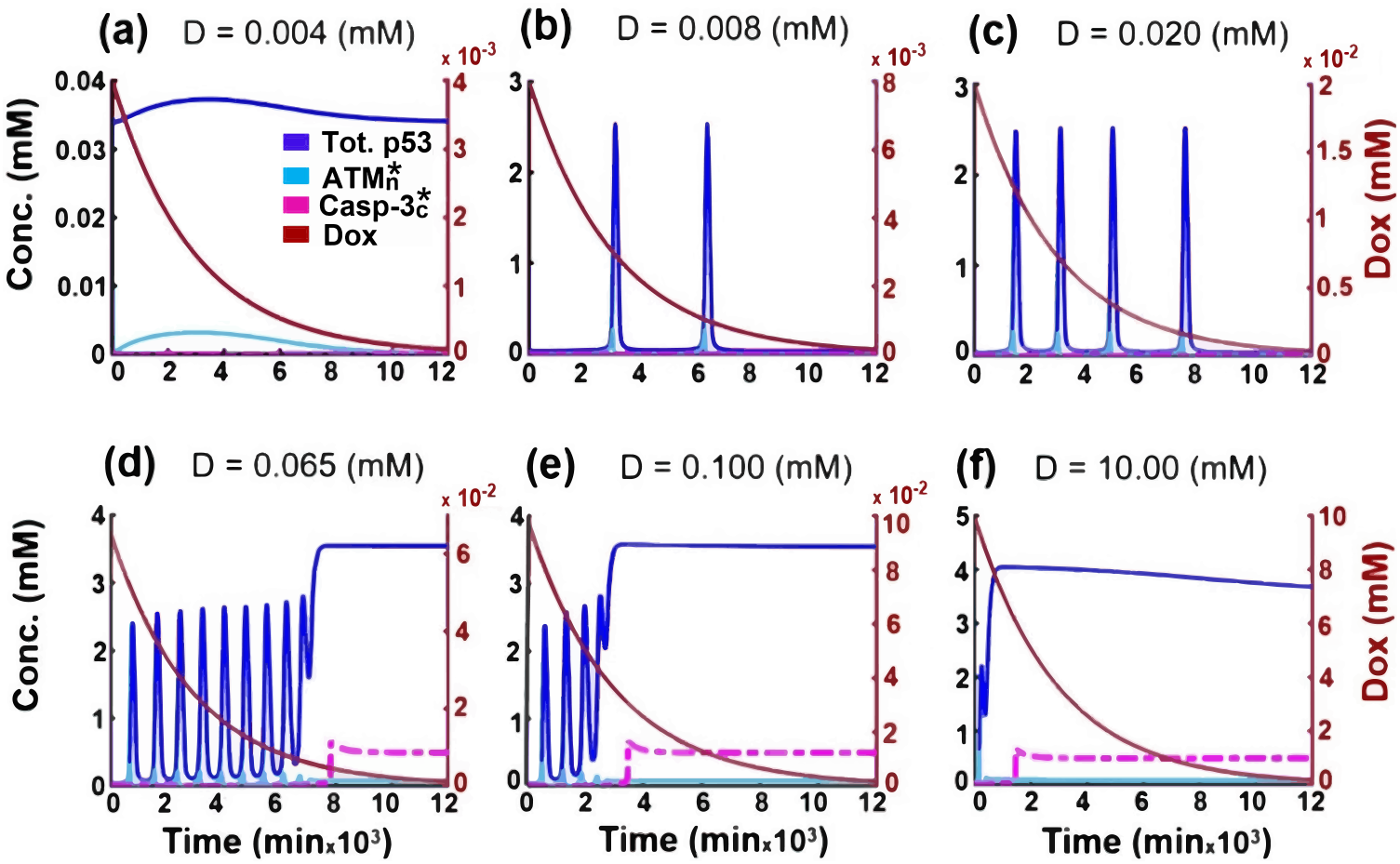
Simulating the cell behaviour after clearing Dox. We set the clearance rate parameter c = 0.00034 (1/min), corresponding to a half-life of 34 hrs. Dox concentrations are shown in the right red axis, while the left black axis shows all other proteins’ concentrations. **a-c** Under low levels of Dox, cells return to their basal condition once Dox gets removed without triggering apoptosis. **(a)** Very low Dox dosage where p53 concentration remains inactive. **(b)** and **(c)** Higher level of Dox leads to high ATM activation triggering p53 oscillatory dynamic; positively correlated with doses. As Dox decays, the proteins’ levels return to basal conditions (indicating survival). **d - f** shows cells trigger apoptosis irreversibly under moderate to high Dox dosages. Here the concentrations remain in the apoptosis steady state even after Dox clearance. **(d)** and **(e)** Under middle Dox levels, cells induce few pulses before irreversibly enacting apoptosis. **(f)** Faster apoptosis decision under acute levels of Dox.

In brief, the system does not remain in the oscillatory regime under low Dox dosages; alternatively, it returns to its normal condition once the drug is cleared. However, under middle and high treatment of Dox, our system settles on the apoptosis steady state and remains there even after clearing Dox from the cell. This finding indicates that apoptosis is an irreversible decision as active caspase-3 remains high and never comes back to a low level. By comparison with experimental data conducted by Wu et al., our model shows a good agreement for cell behaviour under the influence of different Dox levels [4].

### 3.3. Active ATM kinase concentrations govern the transition between p53 dynamics modes

ATM kinase acts as a sensitive and reliable detector of the DSBs induced by the chemotherapy Dox [4, 24, 25]. After sensing the damage, it produces an “on-off” switching signal to the downstream p53 oscillator depending on the amount of DNA damage within the cell. Not only that, but it also promotes the occurrence of the p53 terminal pulse to induce apoptosis if the damage is beyond repair.

In our model, at very low dosages of Dox (*D* < 0.005 mM), the active ATM expression level rises slightly, then quickly returns to its basal level without exciting p53 as it remains predominantly in the inactive form (Fig. 3a). In contrast, a sufficiently strong dose of Dox (0.065 mM > *D* ≥ 0.005 mM) triggers enough ATM activation level, switching on the p53 oscillatory dynamic. This switching behaviour occurs when the activated ATM level exceeds a threshold, driving its protein level excessively high. However, due to negative feedback loops, active ATM concentration repeatedly returns to low levels evaluating the residual damage before inducing the next pulse (Figs. 3b, 3c) and (Fig. 4).

**Figure 4:**
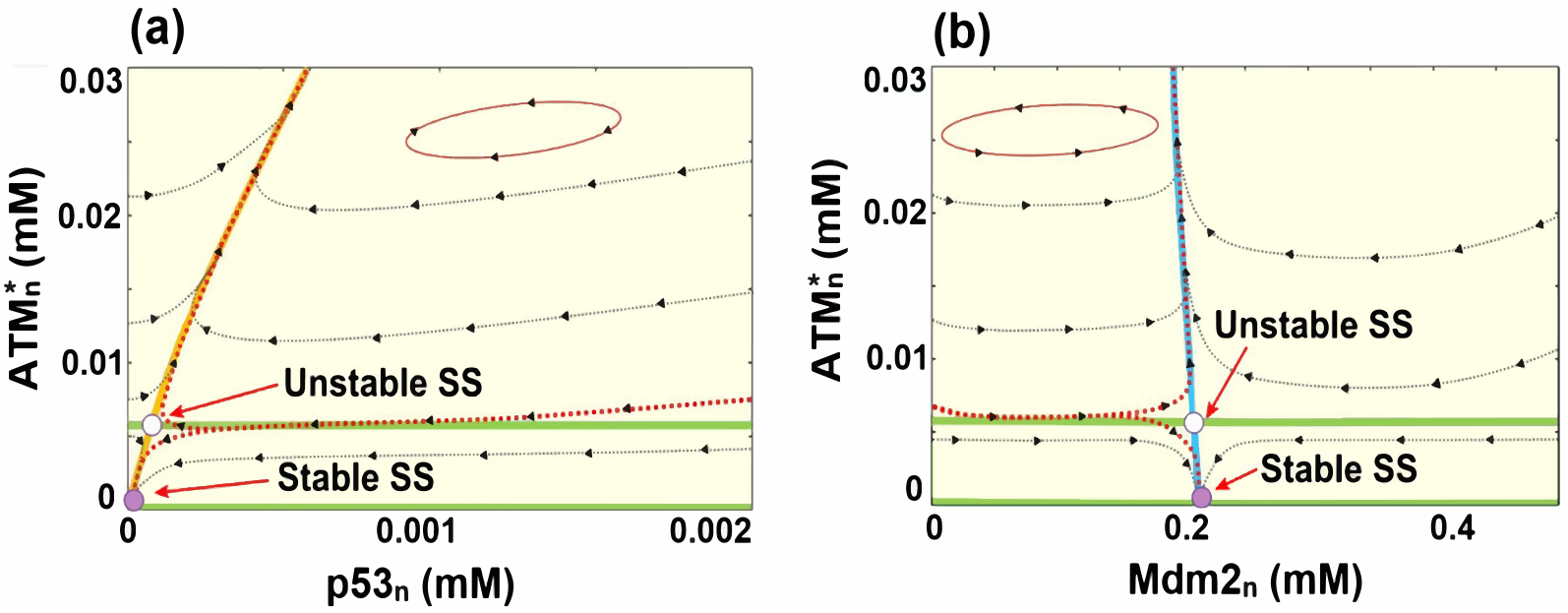
Phase portrait of the system under low Doxorubicin doses (*D* < 0.065 mM). **(a)** Nullcline corresponding to nuclear p53 (p53_*n*_) and active ATM 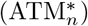. **(b)** Nullcline corresponding to nuclear Mdm2 (Mdm2_*n*_) and active ATM 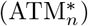. The orange, blue, and green lines represent nuclear p53, nuclear Mdm2, and active ATM nullclines, respectively. The solid magenta dots denote stable equilibria, while the hollow magenta dots denote unstable equilibria. The nullclines demonstrate that elevating 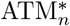 concentrations above the unstable equilibrium will switch on the p53 oscillatory dynamic. Increasing the 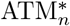 level will promote and degrade the nuclear p53 and Mdm2 levels, respectively. Then, the subsequent negative feedback loops will return 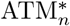 to low concentrations to evaluate the residual damage before inducing the next pulse. So if there were no DNA damage or insufficient Dox to drive 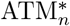 above the unstable equilibrium, the system would switch off the oscillation behaviour heading toward the survival equilibrium.

Moreover, our simulations suggest that the occurrence of the p53 terminal pulses relies heavily on ATM activation levels. Indeed, in response to a moderate dose of Dox (*D* ≥ 0.065 mM), 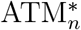 converges to a relatively high constant level in a damped-oscillation manner, followed by a high-amplitude terminal pulse of p53 (Figs. 3d, 3e). Precisely, we observed that as long as 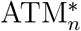 is returning to low levels, just as in the first phase, the cell can still survive, whilst the later phase indicates that the apoptosis decision is made. Specifically, this is when the negative feedback loops fail to significantly lower the 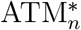 level while it is preparing for the p53 terminal pulse.

### 3.4. The slow rates of Dox clearance enhance its efficacy

Continuing our investigation regarding the effects of Dox clearance and cell state, we gradually decreased the clearance rate parameter *c* in our model. Interestingly, we found that lowering the clearance rate value could ease the occurrence of p53 terminal pulse even under relatively low levels of Dox (Fig. 5). The model results indicate that apoptosis is triggered by exceeding a critical threshold which changes proportionally to the clearance rate under different Dox dosages (Fig. 6). Our finding could be explained by assuming that repeated drug exposure at equivalent concentrations over longer periods may cause a higher number of DSBs, resulting in death fate choice even under low concentrations of Dox. This result agrees with experimental data demonstrating that the longer-lasting presence of Dox or increased Dox bioavailability leads to a significant decrease in tumour size without an increase in Dox concentration [49].

**Figure 5:**
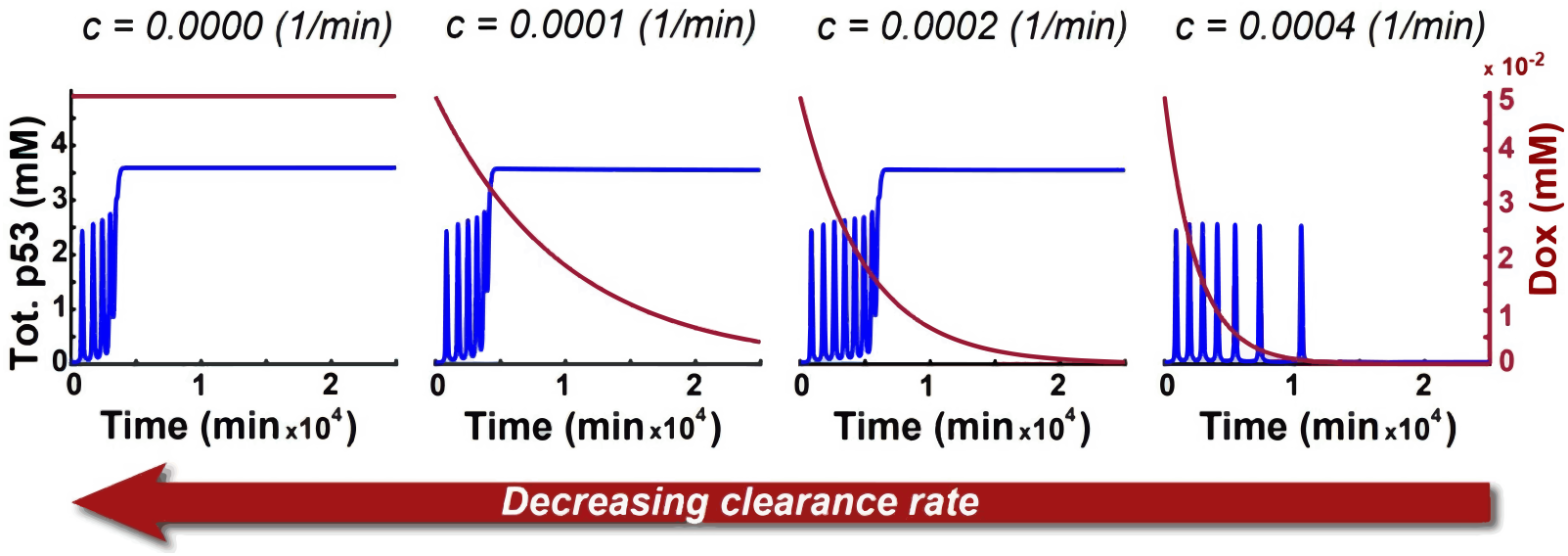
Total p53 concentration under different Dox clearance rates. Under the same concentration of Dox (*D* = 0.05 mM), decreasing the clearance rate (*c*) can ease and accelerate the p53 terminal pulse, thereby promoting cell death.

**Figure 6:**
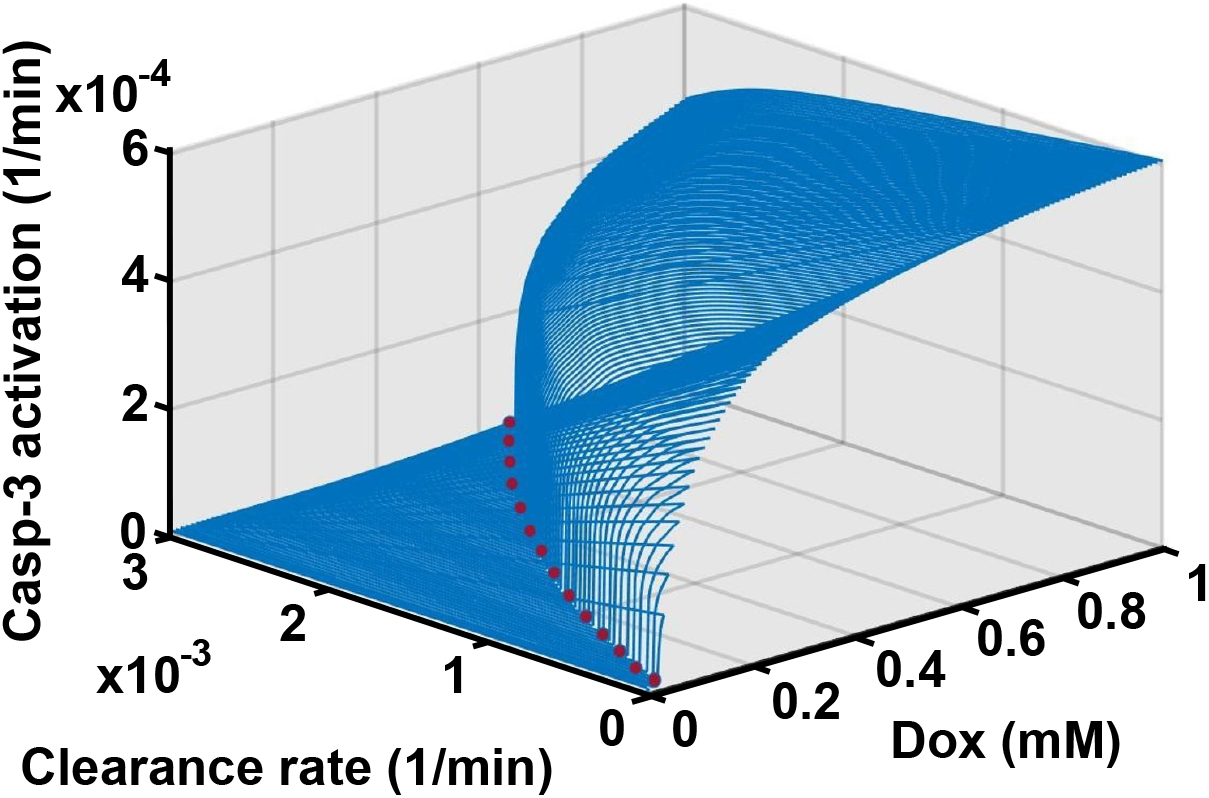
Surface depicting the effect of Dox clearance rate on cellular response to treatment. Here it is possible to observe Dox doses with caspase-3 activation time under different clearance rates where apoptosis can be triggered by crossing a threshold (depicted by the red dotted line).

### 3.5. Two fates for low Dox doses but only one for higher doses

As shown previously, upon Dox treatment, our model’s solutions always tend to one of two stable states with or without the oscillation phase: survival or apoptosis. However, by considering the caspase-3 nullclines, one can see that under low treatment of Dox, the model exhibits three equilibria; two stable and one unstable (Figs. 7a, 7c). At higher Dox doses, however, our model exhibits only one stable equilibrium (Figs. 7b, 7d).

**Figure 7:**
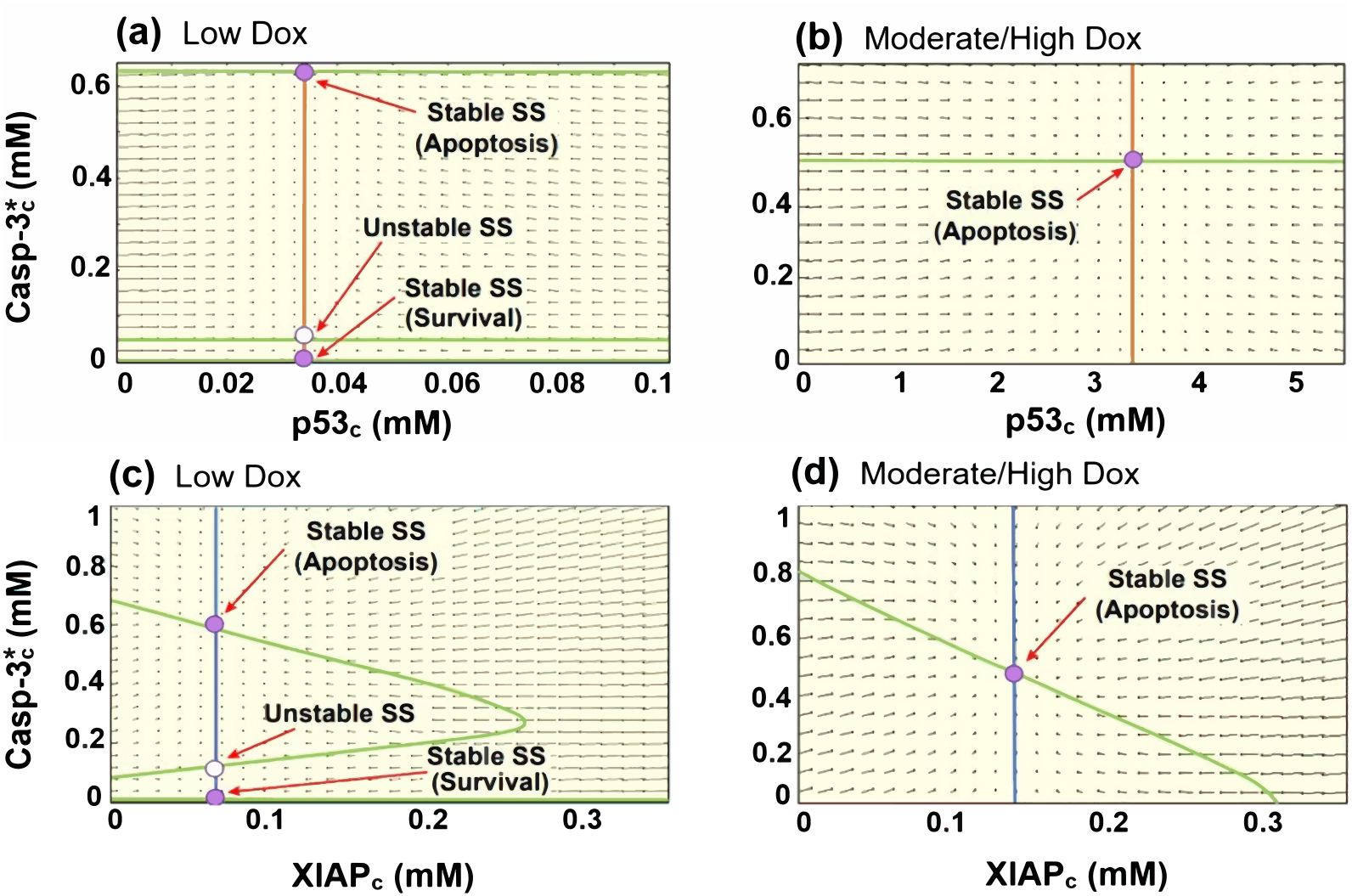
Phase space of the system under different Dox dosages. (**a** and **b**) Nullclines of cytoplasmic p53 (p53_c_) and active caspase-3 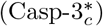. (**c** and **d**) Nullclines of XIAP (XIAP_*c*_) and active caspase-3 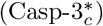. The orange, blue, and green lines represent p53_*c*_, XIAP_*c*_, and 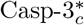 nullclines, respectively. The solid magenta dots denote stable equilibria, while the hollow magenta dots denote unstable equilibria. Under low Dox mode (*D* < 0.065 mM) (**a** and **c**), the system shows two stable equilibria with different concentration levels of active caspase-3. The stable equilibrium with a low concentration of active caspase-3 represents a survival steady state, while the one with a high active caspase-3 concentration represents an apoptosis steady state. Under middle/high Dox mode (*D* ≥ 0.065 mM) (**b** and **d**), the system exhibits a single stable equilibrium (apoptosis steady state) after triggering apoptosis.

By comparing the model’s variables concentration in each steady state under low doses case, we found that these steady states differ only in the caspase-3 concentrations, while the concentrations of the other proteins remain constant at their basal levels. Since caspase-3 is considered a marker of apoptosis, the stable steady state corresponding to a low level of active caspase-3 represents a survival steady state, whereas the one with an elevated active caspase-3 concentration refers to an apoptosis steady state. This result raises the question of under which condition will the cell be driven to the apoptosis steady state if it has no or minor DNA damage? We can answer this question by considering that these steady states differ only in caspase-3 concentrations. Thus, any fluctuation in active caspase-3 concentration due to other stresses (not because of DNA damage) could lead to apoptosis. In particular, if the active caspase-3 concentration solution follows a trajectory above the unstable steady state, the cell will undergo apoptosis regardless of other proteins’ levels within the cell. This finding provides a clue on how much caspase-3 activity may be tolerated due to any accidental fluctuation without the subsequent appearance of apoptotic phenotype.

When subjected to moderate and high drug doses, cells will tend to survive only if the damage gets repaired early with insufficient Dox to cause further damage. Otherwise, the cell will irreversibly undergo apoptosis. The caspase-3 nullclines (Figs. 7b, 7d) were constructed by starting near the apoptosis steady state to show uniqueness stability and apoptosis irreversibility. As expected, they show that after triggering apoptosis by inducing high active caspase-3 levels, repairing the damage or reducing the Dox level cannot cause a transition back from high to low active caspase-3 levels as there is no other stable equilibrium.

### 3.6. Mdm2 expression levels contribute to the p53 response mechanism to treatment, and its amplification leads to apoptotic resistance

Mdm2 amplification has been detected in many human malignancies, and many studies have confirmed its association with chemotherapeutic resistance [50]. Here, we study the effect of Mdm2 overexpression on the system’s dynamical behaviour by assessing the effect of the cytoplasmic Mdm2 basal production rate (*k*_11_) on the p53 response.

According to Fig. 8, under relatively low and high Mdm2_*c*_ basal production rates (*k*_11_), cells always have a unique stable steady state, representing apoptosis and survival steady state, respectively. In contrast, when 0.001 ≤ *k*_11_ < 0.00382, the cell will have to decide whether to survive or die depending on the dose level as it has two stable steady states with high and low p53 concentrations. Our system shows a Hopf bifurcation near *k*_11_ = 0.00382. By crossing this Hopf point to the left, the p53 concentration performs repeated oscillations before settling on one of the equilibria. Furthermore, as *k*_11_ decreases, the number of p53 pulses that occur before achieving the apoptosis steady state decreases while its amplitude increases. However, by keep decreasing the *k*_11_, the oscillation behaviour of p53 disappears, and the cell becomes more and more sensitive to the dose as it heads monotonically toward the apoptotic state under low Dox levels.

**Figure 8:**
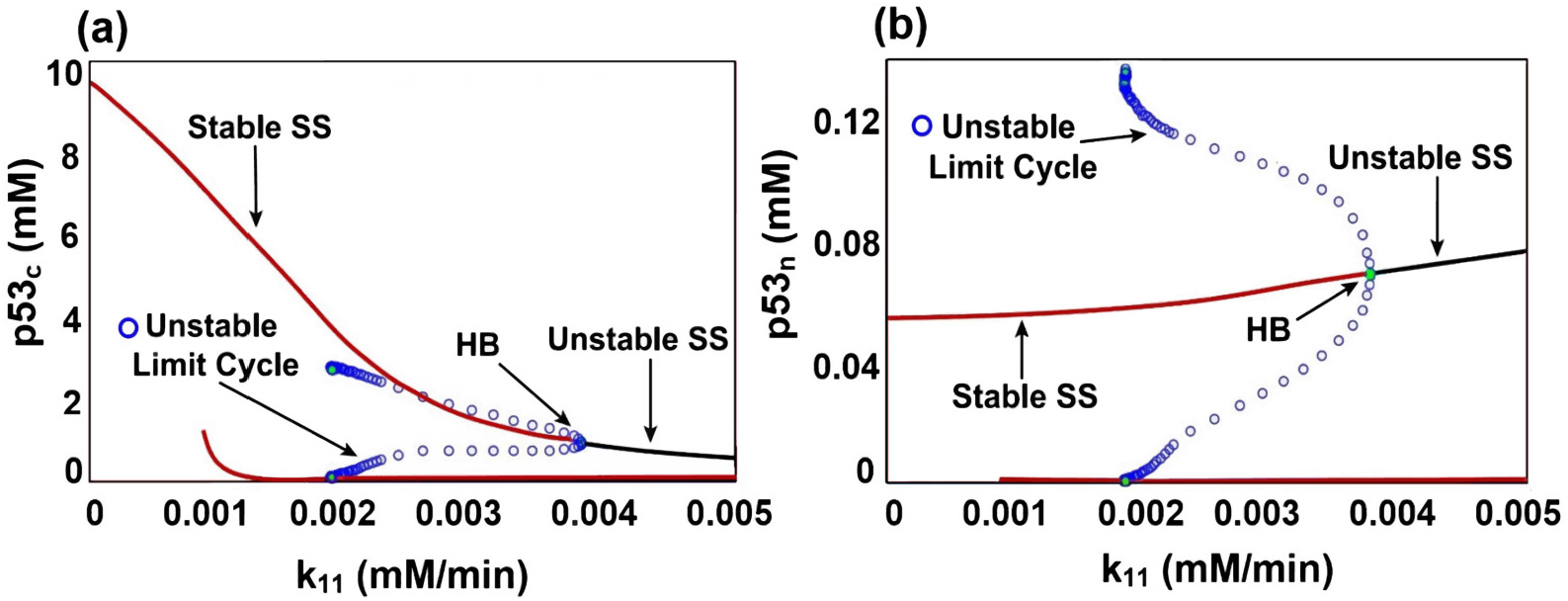
Bifurcation diagrams of p53 level driven by cytoplasmic Mdm2 basal production rate (*k*_11_). **(a)** The cytoplasmic p53 (p53_*c*_) level versus *k*_11_ rate. **(b)** The nuclear p53 (p53_*n*_) level versus *k*_11_ rate. The diagrams show unique stability under high and low *k*_11_, representing survival and apoptosis states, respectively. However, *k*_11_ = 0.00382 represents a Hopf bifurcation point that shifts the system to a bistability regime with two stable steady states (survival and death states). Depending on the dose strength, p53_*c*_ and p53_*n*_ levels settle on one of these steady states after multiple pulses. The amplitude of these pulses increases as *k*_11_ decreases, but at some point, the cell becomes highly sensitive to the treatment, heading monotonically toward the apoptotic state without oscillating even under minor damage.

In short, our simulation shows that low expression levels of Mdm2 enhance the cells’ sensitivity to chemotherapy and facilitate the apoptotic program. In contrast, cells with abnormally high Mdm2 basal transcription rates would resist treatment even under high doses. Furthermore, the Mdm2 expression level is seemed to play a role in generating the p53 oscillatory behaviour and shaping its amplitude.

Several studies asserted that targeting the E3 ubiquitin ligase activity of Mdm2 by using an Mdm2 inhibitor such as Nutlin-3 would increase the sensitivity of tumour cells to radio- or chemotherapy [51, 52, 53]. Nutlin-3 is a widely studied small molecular inhibitor of Mdm2, which disrupts the p53-Mdm2 interaction by binding the N-terminus of Mdm2 (p53 binding site) [54]. To further validate our model, we examined the effect of Nutlin-3 on the p53 cellular level by decreasing the Mdm2_*c*_-dependent p53_*c*_ degradation rate (*k*_8_) (Fig. 9). Accordingly, we found that silencing the Mdm2 in combination with chemotherapy Dox increases the accumulation levels of p53 and accelerates the induction of p53 terminal pulses. This result coincides with experimental observations that show promoted accumulation of p53 and early onset of p53 terminal pulse with a significantly increased apoptotic rate in response to Dox treatment followed by Nutlin-3 [4].

**Figure 9:**
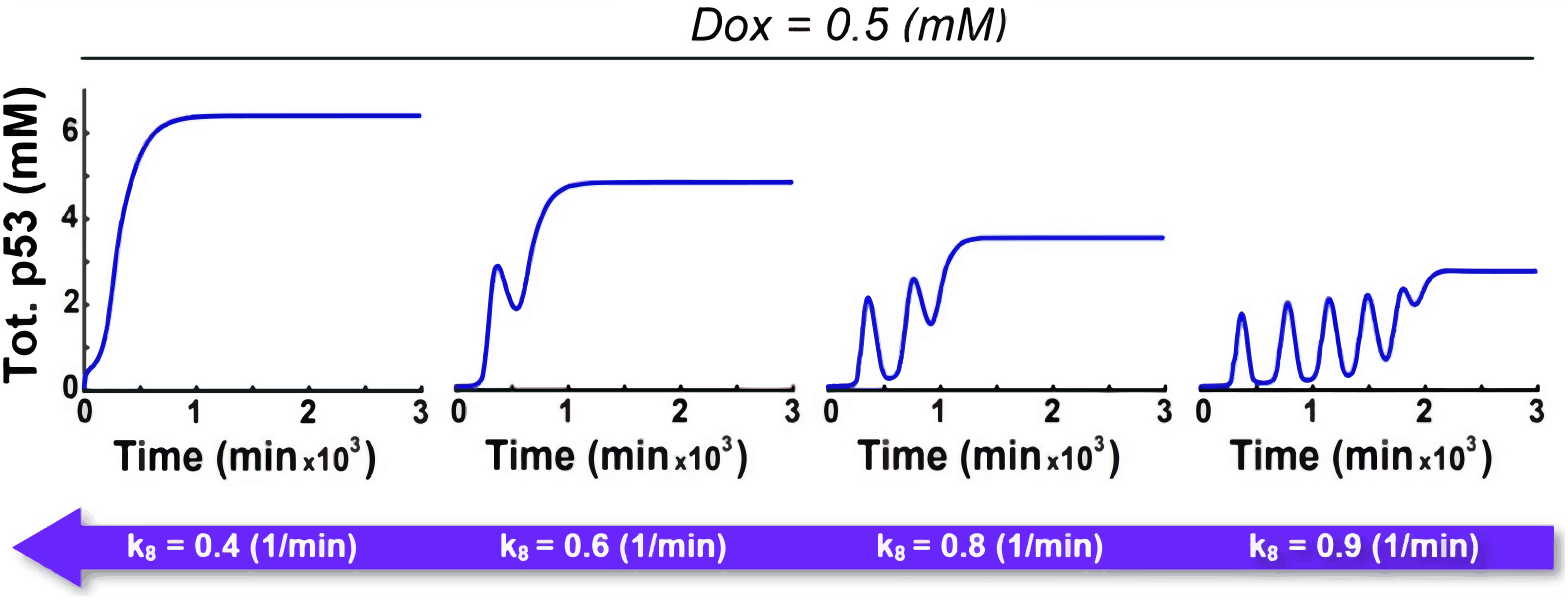
Simulation reproduces the effect of Nutlin-3. The figure showcases the total p53 protein concentration stimulated by Dox = 0.5 mM under different Mdm2_*c*_-dependent p53_*c*_ degradation rates (*k*_8_). Decreasing the Mdm2_*c*_-dependent p53_*c*_ degradation rate (*k*_8_) boosts the accumulation of p53 and accelerates the generation of p53 terminal pulses.

### 3.7. Upregulation of XIAP by Mdm2 following chemotherapy attenuates the apoptotic pathway subsequent to MOMP

Mdm2 can play several p53-independent roles in cancer cells, as it regulates the XIAP expression level following radio- and chemotherapy, allowing for a rapid response when XIAP function is required [5, 15, 43]. Mdm2 interacts with the XIAP mRNA and activates the IRES-dependent XIAP translation during cellular stress, resulting in overexpression of XIAP [45]. This upregulation of XIAP was found to contribute enormously to the drug resistance of tumour cells [5]. Therefore, here we aim to capture the principle of how the upregulation of XIAP can attenuate the apoptotic pathway after MOMP. To do so, we investigated the effect of the XIAP induction rates (*k*_24_) on the total p53, XIAP, and active caspase-3 concentrations under fixed Dox doses (*D* = 0.1 and 1 mM) (Fig. 10).

**Figure 10:**
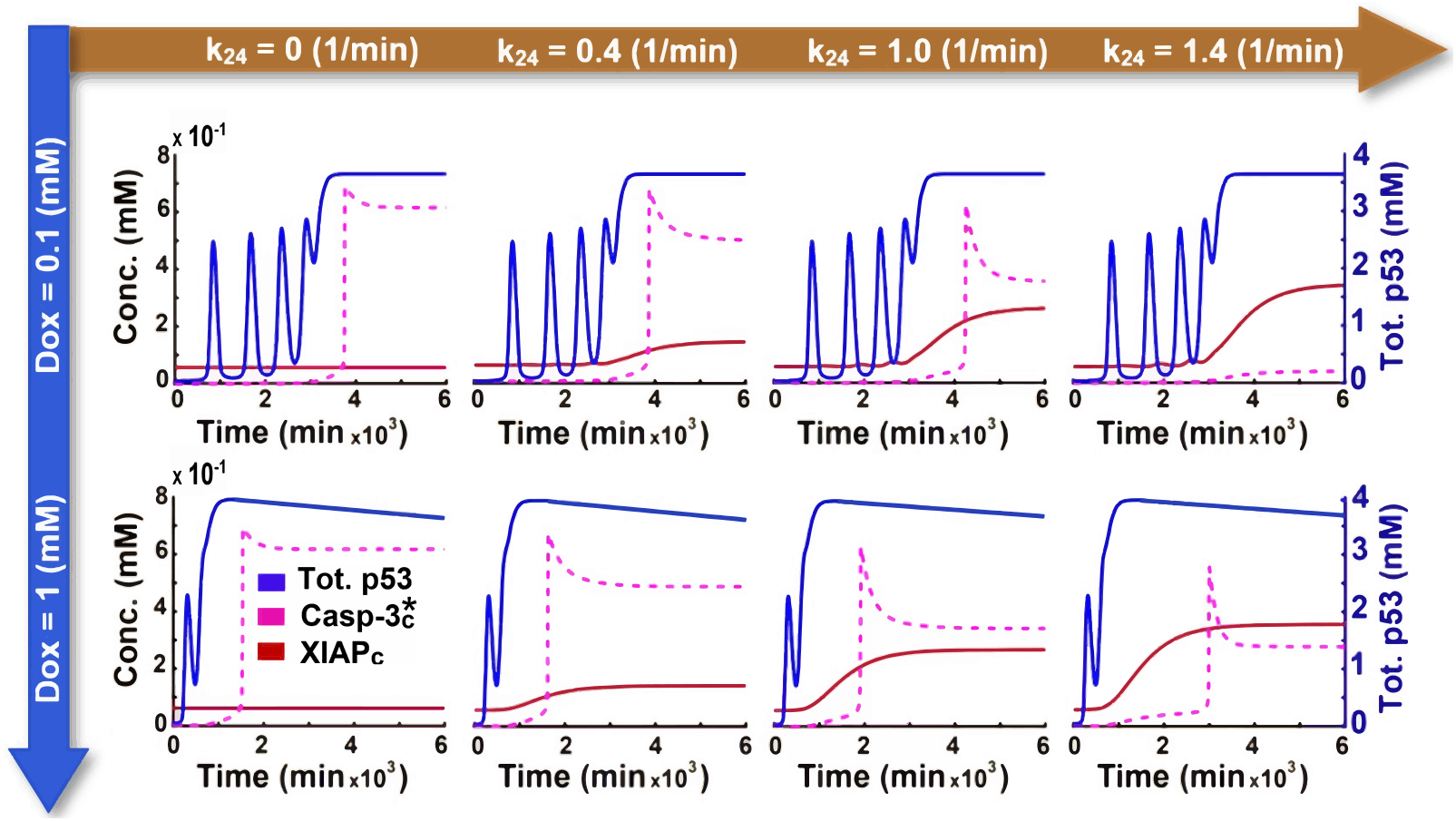
Simulating the effect of XIAP induction rate (*k*_24_) on the cell fate. It shows the time course of the total p53 (right axis), XIAP (XIAP_*c*_), and active caspase-3 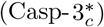 (left axis) concentrations stimulated by a moderate dose (Dox = 0.1 mM) and high dose (Dox = 1 mM) under different XIAP induction rates (*k*_24_). Increasing the XIAP induction rate (*k*_24_) leads to low caspase-3 activation levels and delays in their activation. However, beyond *k*_24_ = 1.4, the caspase-3 activity can entirely block under middle Dox doses, specifically when the XIAP_*c*_ level exceeds 0.34 mM.

The simulation results indicate that increasing the XIAP induction rate boosts the XIAP concentration level, leading to a significant delay in caspase-3 activation. Moreover, active caspase-3 is settling on a lower level, suggesting an even larger delay in triggering apoptosis. Additionally, pushing the XIAP level further (*X_c_* > 0.34 mM) may completely block caspase-3 activation under middle-Dox doses. Our findings are consistent with experimental data that overexpression of XIAP slowed down the effector caspase activation and significantly blocked the substrate cleavage when its concentration exceeded 0.30 mM [55].

Moreover, our results show that p53 ultimately tend to reach the same concentrations under all moderate to high doses of Dox regardless of XIAP levels. In contrast, caspase-3 may trigger or not depending on the XIAP induction rate, determining whether the cell will die or resist (Fig. 10). In other words, the failure to activate apoptosis in some cancer cells with wild-type p53 might be due to heterogeneity between cells in upregulating the XIAP protein rather than the p53 protein level. Furthermore, under relatively high XIAP induction rates, cells treated with high doses of Dox accumulate p53 faster and manage to trigger apoptosis, while cells with moderate Dox levels do not (Fig. 10). These results show a complete agreement with Paek et al., which reported that apoptotic cells did not have a higher maximum of p53 levels but did accumulate p53 earlier than surviving cells [5].

### 3.8. Disrupting the XIAP-caspase-3 or XIAP mRNA-Mdm2 interaction improves Doxorubicin efficacy

Several therapeutic strategies have been designed to target XIAP protein and overcome chemotherapy resistance. One strategy suggests targeting the XIAP-caspase interaction, disabling XIAP from inhibiting caspases by using a small molecule called LCL-161. This molecule simply binds the common BIR domain of XIAP, where it binds caspase-3 and -9 [5, 56].

Here, we investigate the influence of XIAP inhibitor on the caspase-3 activation by constructing a bifurcation diagram showing the active caspase-3 concentration levels (when *D* ≥ 0.065 mM) versus the XIAP_*c*_-dependent 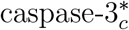 degradation rate (*k*_30_) (Fig. 11a). Notably, minimising the XIAP function helps activate a high level of caspase-3, thereby inducing apoptosis more easily, which is in good agreement with experiments that showed a decreased cell viability following Cisplatin treatment combined with LCL-161 [5]. By contrast, elevating the interaction rate *k*_30_ shifts the cells to a bistability regime with two stable steady states corresponding to survival and apoptosis. However, when *k*_30_ increases beyond 0.055 per min, which can be due to Smac scarcity, the cell will resist and survive as it has only one stable steady state accompanied by a low concentration of active caspase-3.

**Figure 11:**
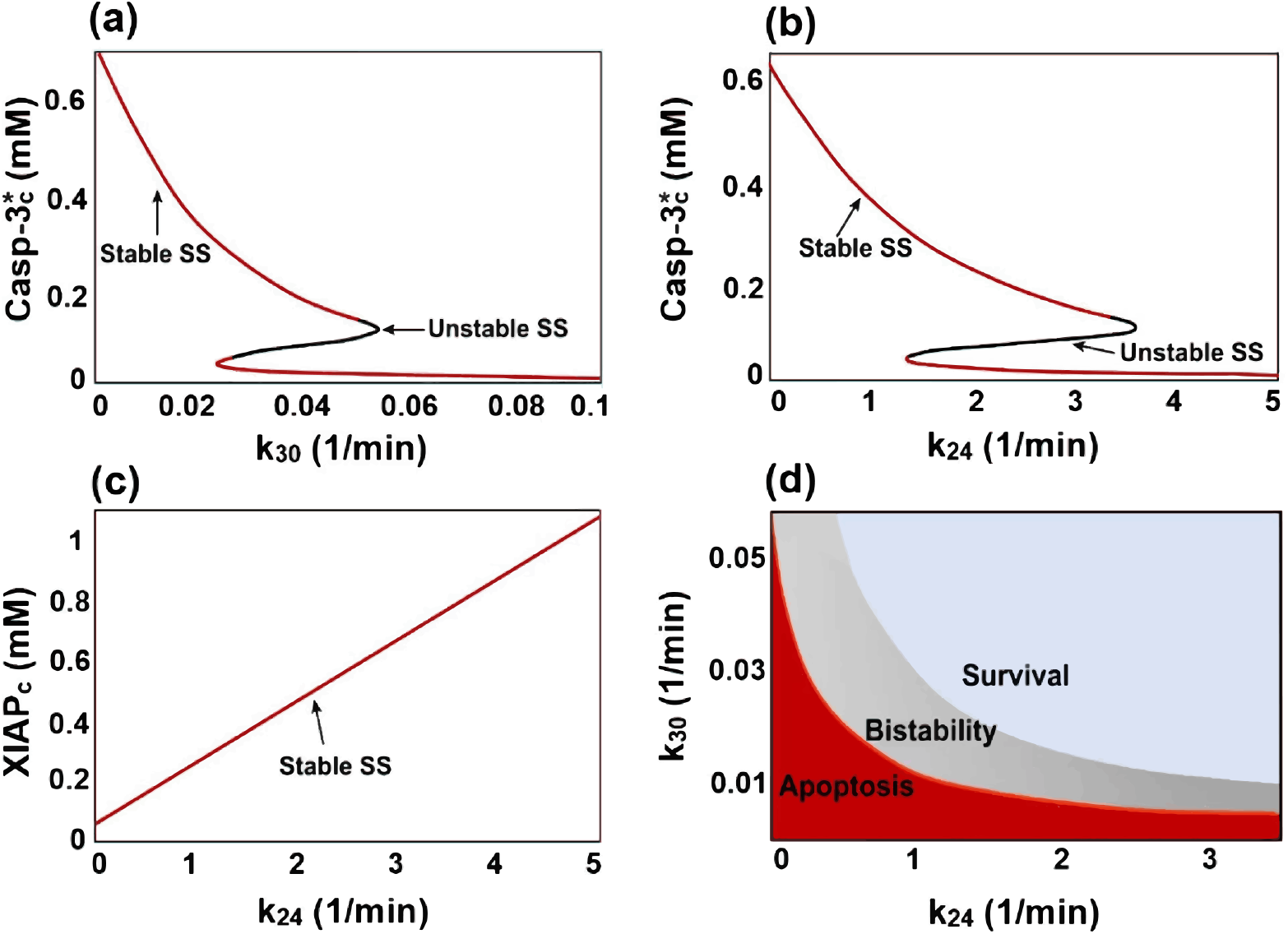
Bifurcation diagrams illustrating the effect of the degradation rate of active caspase-3_*c*_ by XIAP_*c*_ (*k*_30_) and the XIAP_*c*_ induction rate by Mdm2_*c*_ (*k*_24_) on the cellular response. **(a)** Bifurcation diagram of 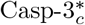 level in the apoptotic steady state vs degradation rate of 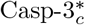 by XIAP_*c*_ (*k*_30_). Under low *k*_30_ rates, the model readily enacts apoptosis by activating enough caspase-3. The system exhibits two stable equilibria (survival and apoptosis) when 0.025 < *k*_30_ < 0.055. Increasing the value of *k*_30_ beyond 0.055 per min would allow the cell to survive and resist Dox. **(b)** and **(c)** Bifurcation diagrams of 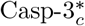 and XIAP_*c*_ levels in apoptosis steady state vs XIAP_*c*_ induction rate by cytoplasmic Mdm2 (*k*_24_). Suppressing the induction of XIAP facilitates executing apoptosis by triggering a high 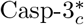 level. Nevertheless, crossing the bifurcation point *k*_24_ = 1.4 elevates the XIAP_*c*_ concentration above 0.34 mM, instigating a bistability regime with two stable equilibria corresponding to apoptosis and survival. However, increasing the induction rate further more (*k*_24_ > 3.4) returns the system to a unique stable equilibrium with low concentrations of 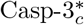. This means apoptosis may be completely blocked even under high concentrations of Dox. **(d)**. Cell fate map represented by a two-parameter bifurcation diagram for *k*_30_ and *k*_24_, where apoptosis, bistability, and survival regions are shaded in red, grey, and blue, respectively.

On the other hand, recently, a new approach has been taken to overcome the XIAP upregulation by using a novel Mdm2/XIAP dual inhibitor called MX69 [57, 58]. It is shown to have a robust cytotoxic effect in tumour cells by blocking the physical interaction between Mdm2 and XIAP mRNA, inhibiting IRES-dependent translation of XIAP [57, 58]. To further validate our model, we examined the effect of the inhibitor MX69 on the cell outcomes by displaying the levels of active caspase-3 and XIAP driven by the XIAP induction strength (*k*_24_) when *D* ≥ 0.065 mM (Figs. 11b, 11c). As expected, suppressing the induction of XIAP facilitates executing apoptosis once irreparable damage is detected. However, the bistability regime appears by crossing the bifurcation point around *k*_24_ = 1.4, strictly when the XIAP level exceeds 0.34 mM. In this case, acute doses of Dox may activate the death fate if the solution of active caspase-3 heads above the unstable equilibrium; otherwise, the cell will resist and escape apoptosis due to incomplete substrate cleavage under low levels of active caspase-3. Furthermore, the model predicts that apoptosis may be entirely blocked even under high concentrations of Dox when the induction rate *k*_24_ exceeds 3.4 per min, which is responsible for increasing the XIAP level more than fourteen fold higher than its physiological concentration.

Our analysis has shown how regulating the XIAP intracellular levels and its ability to repress caspases activation play a crucial role in therapeutic response. Therefore, it would be beneficial to get a comprehensive view of their impacts simultaneously on cell fate. To do so, we constructed a two-parameters bifurcation diagram as a cell fate map determined by the parameters: *k*_24_ and *k*_30_ (Fig. 11d). The cell fate map is divided into three regions (delimited by the two bifurcation curves in Figs. 11a, 11b): apoptosis, survival, and bistability representing high caspase-3 activation, low caspase-3 activation, and moderate/low caspase-3 activation regions, respectively.

## 4. Discussion

Cancer cells develop resistance to almost all anticancer drugs via a variety of distinct mechanisms and pathways. Agents commonly used in cancer chemotherapy often rely on the induction of cell death via the p53 tumour suppressor gene. The mathematical model proposed in this paper provided detailed insight into the complex nonlinear signalling processes controlled by p53 in the apoptosis pathway in response to chemotherapy Doxorubicin. Results demonstrate that the variation in p53 dynamics between individual cells can be responsible for different responses to chemotherapy. p53 oscillations that do not reach a critical plateau of terminal pulse fail to trigger MOMP, resulting in a lack of response to chemotherapy. Thus, the p53 terminal pulse is considered an important sign of disrupting mitochondrial membrane potential as an essential step toward executing apoptosis.

One of the most common disorders seen in cancer cells believed to contribute significantly to chemotherapy resistance is Mdm2 amplification [50]. Our model demonstrated how Mdm2 overexpression could be involved in the p53 dynamics variability between cancer cells. Despite this vital role of Mdm2 in controlling the p53 dynamical responses, most studies have ignored their ability to regulate the XIAP protein at the translational level [45]. Our study emphasises the importance of the anti-apoptotic mechanisms of XIAP, as excessive stimulation of XIAP by Mdm2 following chemotherapy may lead to evading apoptosis even after the MOMP.

Previous studies have shown some aspects of the p53 dynamics in response to DNA damage under various stimuli [59, 60, 61, 62], but none of them has shown all three modes of p53 in response to different stimulus strength that has been observed experimentally [4]. p53 can switch between three different modes depending on the DNA damage/dose levels ranging from oscillation only, oscillation followed by a high-amplitude terminal pulse, and monotonic increase. A recent study by Sun et al. has modelled the dual-phase p53 dynamics under Doxorubicin treatment based on the contested “affinity model” without considering the effect of XIAP overexpression on cellular outcomes [63]. In their model, the occurrence of the p53 terminal pulse can be considered an ultimate measure of cell fate. However, that contrasts with Peak et al.’s experimental observations that showed no difference in the maximum p53 protein levels between some apoptotic and survival cells with a remarkable increase of anti-apoptotic proteins, such as XIAP, in resistant cells [5]. Therefore, Sun et al.’s model cannot be applied to infer the cellular responses of cells that experience an increase in XIAP following treatment.

In addition, our model demonstrated the irreversibility of the apoptosis decision by incorporating the Dox clearance mechanisms, showing the actual ultimate cell fate after eliminating Dox. Furthermore, our study provides preliminary information on how the Dox clearance rate may affect therapeutic resistance and reduce the level of doses required to induce cell death.

Despite our model’s success in reproducing and explaining experimental observations, it has some limitations. Our current work focused only on the intrinsic pathway in activating apoptosis. However, the extrinsic pathway may trigger under acute chemotherapy doses by producing high reactive oxygen species (ROS) levels [64]. ROS are generated after exposure to physical agents (ultraviolet rays and heat) and after chemo- and radiotherapy in cancer. ROS enhance the assembly of DISCs, thereby triggering the extrinsic apoptotic pathway. Additionally, it can improve the intrinsic pathway activity by increasing the ubiquitination of the anti-apoptotic protein Bcl-2, disrupting mitochondrial membrane potential [64]. This would explain the activation of apoptosis and overcome the high induction of XIAP under severe chemotherapy dosages. The model presented here could be extended to incorporate ROS signalling networks so that we could gain more accurate predictions of the caspases activation levels and the cell fate in response to chemotherapy.

Some studies have confirmed an increase in other members of the IAP family, such as cIAP1, cIAP2, and ML-IAP, increasing the viability of cancer cells following chemotherapy treatment [1, 5]. Increased IAP expression inhibits two separate apoptotic pathways differentiated by their dependence on caspase-8. cIAP1 and cIAP2 expression have been seen to prevent caspase-8-dependent apoptosis, while ML-IAP and XIAP expression inhibits caspase-8-independent apoptosis [5]. For simplicity, the current model considers only the upregulation of XIAP in response to chemotherapy. However, adding other signalling networks, such as ROS, may require incorporating the other IAP members that influence the extrinsic apoptosis pathway (caspase-8-dependent apoptosis).

Recently, tumour metabolism has been seen to be also incorporated into chemoresistance [65]. Unlike normal cells, tumour cells preferentially metabolise glucose by glycolysis instead of oxidative phosphorylation (OXPHOS), even in the presence of oxygen. The so-called “Warburg effect” is a critical step in cancer development and contributes to chemoresistance, but the underlying mechanisms of action remain unclear [65]. Incorporating other organelles, such as mitochondria and the signalling pathway in the tumour metabolism would give a more comprehensive picture of the resistance mechanisms.

In conclusion, our model offers a broader understanding of the mechanism of chemotherapy resistance. We investigated the p53 dynamics under various Doxorubicin dosages and examined different scenarios that might contribute to resistance. Importantly, this study modelled for the first time the observations of the fractional killing of cancer cells by incorporating the effect of XIAP upregulation rates following chemotherapy. Our study suggests that modifying p53 protein dynamics and targeting the induction of XIAP by Mdm2 may offer a novel therapeutic opportunity for designing efficient drug combinations. Additionally, slowing the Dox clearance rate would further improve cellular outcomes and overcome resistance.

## 5. Acknowledgements

The authors would like to acknowledge funding to R.A. from [The Saudi Arabian Cultural Bureau (SACB)]. Cancer Research UK funding to D.T. and E.V.-S. (C42109/A26982 and C42109/A24747). F.S. was supported by a UKRI Future Leaders Fellowship, grant no. (MR/T043571/1). Finally, we would like to acknowledge the support and resources of the Birmingham Metabolic Tracer Analysis Core (MTAC).

## 6. Contribution

**R.A.** conceived the project and the mathematical model, performed in-silico experimentation and analysis, and wrote the manuscript. **E.V.-S.** provided critical technical advice concerning the in-silico modelling, its numerical solution, analysis, and wrote the manuscript. **D.T.** provided critical technical advice on the biology discussed in this article, as well as advice regarding the validity of the model’s results. **F.S.** provided critical technical advice concerning the in-silico modelling and analysis, organised, supervised and managed the study.

## Appendix A. Detailed Model

### Appendix A.1. Model Description

In response to DNA double-strand breaks (DSBs) induced by chemotherapy Doxorubicin (Dox), the ataxia-telangiectasia mutated (ATM) is activated through phosphorylation, which then phosphorylates both p53 and murine double minute 2 (Mdm2), affecting their stability. Let the concentration of active nuclear ATM be denoted by 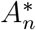. Then,

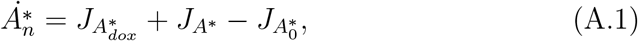

where,

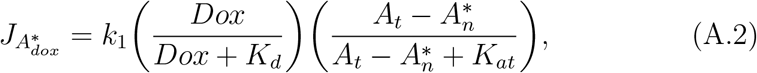

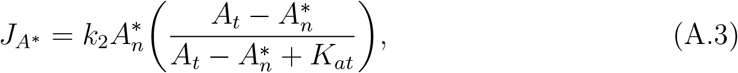

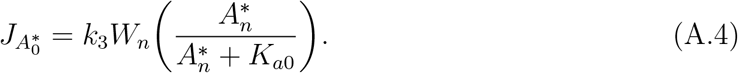

In Eq. (A.2), the phosphorylation rate of ATM by Dox is represented by *k*_1_ where, the phosphorylation of ATM is increased dose-dependently [4]. The flux submodel in this term is based on that of Sun et al., but we added a saturating kinetic [63]. Since the intermolecular ATM autophosphorylation is required for the dissociation of the ATM dimer after DNA damage, we considered positive feedback in which ATM is activated by itself denoted *k*_2_ to be the ATM autophosphorylation rate as in Eq. (A.3), [26]. Note that the total concentration of ATM (*A_t_*) is assumed to be a constant in Eqs. (A.2) and (A.3) because experiments showed that the level of ATM does not change significantly in response to DNA damage [7, 66]. On the other hand, 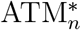 can be inactivated by wild-type p53-induced phosphatase 1 (Wip1) through dephosphorylation, enclosing a negative feedback loop, represented in Eq. (A.4), [13]. Here, *k*_3_ refers to the dephosphorylation rate of 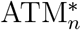 by Wip1; denoted by *W_n_*. Finally, as the phosphorylation and dephosphorylation processes can be considered enzyme-catalysed reactions, they were assumed to follow Michaelis-Menten kinetics in our model [61].

Let *P_c_* and *P_n_* denote the concentrations of p53 protein in the cytoplasm and the nuclear compartments, respectively. Then,

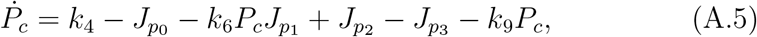

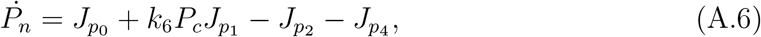

where,

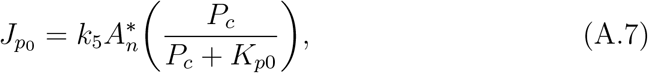

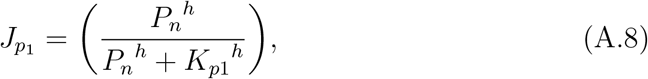

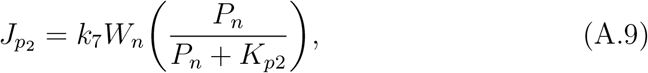

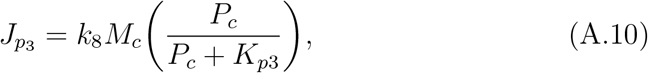

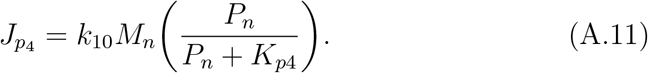

In Eq. (A.5), *k*_4_ represents the basal production rate of p53 in the cytoplasm. However, we assume that p53 will be restricted in the cytoplasm until it gets activated by ATM. Therefore, in Eq. (A.7), the parameter *k*_5_ represents the activation rate of p53 by 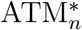 which results in the accumulation of p53 in the nucleus in a tetramer form [4, 27, 32, 33]. Since activated p53 can promote its own activation and nuclear accumulation by inducing phosphatase and tensin homolog (PTEN) [35, 38], we added a term accounting for this positive feedback loop where *k*_6_ is considered as the strength of the feedback loop (Eq. (A.5) and Eq. (A.6)). However, the activation of this positive feedback loop is assumed to be proportional to the nuclear p53 concentration due to its ability to act as a transcription factor, thereby producing PTEN. Moreover, by considering the tetramer form of nuclear p53 while transcribing genes, this term is characterized by a Hill function with a Hill coefficient four (Eq. A.8), [67]. When it comes to Eq. (A.9), *k*_7_ corresponds to the dephosphorylation rate of nuclear p53 catalyzed by Wip1, which enables Mdm2, represented by (*M_c_*) and (*M_n_*) in Eqs. (A.10) and (A.11), to bind p53 and turns on p53 nuclear export by unmasking p53 nuclear export signals (NESs) [13, 29, 34]. Thus, in our model, unphosphorylated nuclear p53 is assumed to be moved into the cytoplasm. On the other hand, the degradation rate of p53 is composed of a basal degradation rate (*k*_9_ - Eq. (A.5)) and an Mdm2-dependent degradation rate (Eqs. (A.10) and (A.11)), [21, 22]. Since Mdm2 shows a weaker binding to nuclear p53 than cytoplasmic p53 [7, 8], we assumed that Mdm2-dependent degradation rate of nuclear p53 (*k*_10_ - Eq. (A.11)) is much smaller than that of cytoplasmic p53 (*k*_8_ - Eq. (A.10)). Furthermore, the degradation of both cytoplasmic and nuclear p53 by Mdm2 was described by Michaelis-Menten kinetics due to the similarity between the catalytic role of Mdm2 as a ubiquitin ligase and the enzyme role in converting a substrate into a product [59].

As stated above, let *M_c_* and *M_n_* denote the concentrations of cytoplasmic and nuclear Mdm2, respectively. Then,

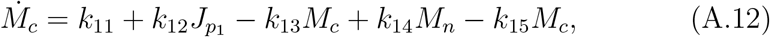

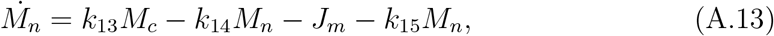

where,

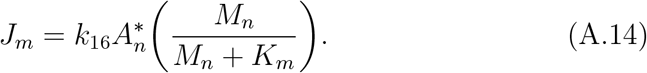

In Eq. (A.12), the parameter *k*_11_ represents the basal production rate of Mdm2 in the cytoplasm, while *k*_12_ accounts for the Mdm2 synthesis by p53 [22, 23], which is controlled by the active tetrameric nuclear p53 and modelled by a Hill function with coefficient four (Eq. (A.8)). On the other hand, Mdm2 is subject to signals that promote its translocation from the cytoplasm into the nucleus [35, 36], and it is exported to the cytoplasm while binding the p53 to mediate the p53 degradation machinery there [29]. Therefore, for the sake of simplicity, we assumed constant rates *k*_13_ and *k*_14_ (Eqs. (A.12) and (A.13)) corresponding to Mdm2 migration from the cytoplasm to the nucleus and from the nucleus to the cytoplasm, respectively. When it comes to the degradation rate, although Mdm2 has a basal degradation rate represented by *k*_15_ (Eqs. (A.12) and (A.13)), its degradation is further stimulated in the presence of active ATM [27]. Consequently, as in Eq. (A.14), we added a degradation rate for nuclear Mdm2 by 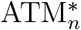 represented by *k*_16_.

Let us denote the concentrations of nuclear Wip1 and active cytoplasmic Bax by *W_n_* and 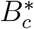, respectively. Then,

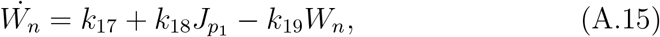

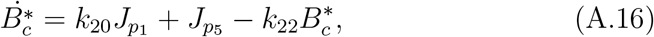

where,

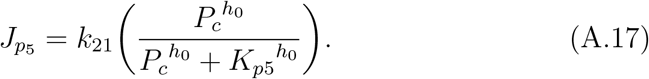

In Eq. (A.15), we represented the basal production rate of Wip1 by a constant rate *k*_17_. Then, we added the contribution of nuclear p53 to the transcription of the Wip1 and active Bax genes (Eqs. (A.15) and (A.16)), [37, 68]. The production rates of Wip1 (*k*_18_) and Bax (*k*_20_) by p53 are modeled by a Hill function as in Eq. (A.8). Although the translation of mRNAs into proteins occurs in the cytoplasm by binding ribosomes, we assumed that once mRNAs of Wip1 are translated into proteins, they are directly moved to the nucleus since Wip1 is predominantly nuclear protein [37]. Furthermore, it has been shown that p53 has transcriptional and non-transcriptional functions to activate pro-apoptotic proteins. Indeed, p53 in the cytoplasm have seen to bind to Bcl-x and Bcl-2, allowing for Bax activation by releasing it from its inhibitors [68, 69, 70]. As a result, we took into consideration the activation rate of Bax by cytoplasmic p53 (*k*_21_ - Eq. (A.17)), which is regulated by a Hill function that gradually increases with the cytoplasmic p53 concentration. Finally, the basal degradation rates for both Wip1 and active Bax are represented by *k*_19_ and *k*_22_ in Eqs. (A.15) and (A.16), respectively.

The concentration of cytoplasmic X-linked inhibitor of apoptosis (XIAP), *X_c_*, changes with respect to time according to,

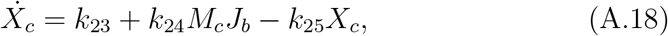

where,

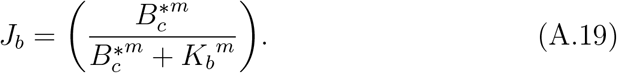

In Eq. (A.18), the basal production and degradation rates of XIAP are determined by *k*_23_ and *k*_25_, respectively. The XIAP expression levels were seen to be increased following chemotherapy drugs such as Cisplatin, Doxorubicin, Etoposide, and Camptothecin [5]. This upregulation of XIAP protein is assumed to be regulated by Mdm2 at the translational level in response to apoptotic stimuli [45], but how Mdm2 are assigned to their translational function is unknown yet [44]. Therefore, we assumed that the induction rate of XIAP (*k*_24_ - Eq. (A.18)) after Doxorubicin treatment is regulated by the cytoplasmic Mdm2, which can interact with the XIAP mRNA in the cytoplasm. However, this term is triggered only when the active Bax level is high enough for mitochondrial outer membrane permeabilization as a sign of apoptotic stimulus. It is modelled by a Hill function that increases with 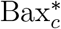 levels but with a relatively high Hill coefficient (*m*) that guarantees the XIAP induction occurs only under apoptotic conditions (Eq. (A.19)).

Now, let the inactive and active concentrations of cytoplasmic caspase-3 be denoted by *C*_3_*c*__ and 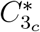, respectively. Then,

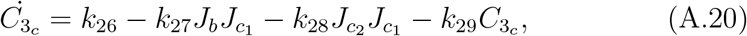

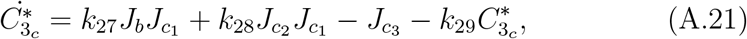

where,

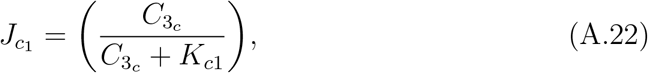

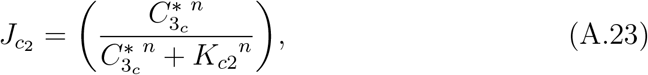

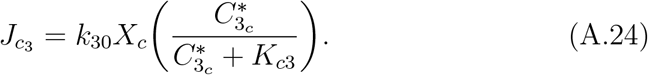

In Eqs. (A.20) and (A.21), we assumed that caspases-3 are always present in inactive form with basal production rate *k*_26_. However, under stress conditions, caspase-3 activation occurs at a rate *k*_27_ modulated by active Bax protein level that has the potential for causing mitochondrial outer membrane permeabilisation (MOMP) [18]. This term comprises several steps and molecular reactions that subsequently lead to caspase-3 activation. Therefore, it is represented here by a Hill function that gradually increases from zero to one with active Bax concentration, where *K_b_*, the Half maximal concentration of the function, represents the inhibitory mechanisms mediated by anti-apoptotic proteins Bcl-2 and Bcl-x (not explicitly included in our model). The Hill coefficient *m* was selected to be high enough to ensure no caspase-3 activation occurs under low levels of 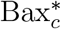 protein as well as no XIAP induction by Mdm2 as in Eq. (A.19). Moreover, since the conversion of caspase-3 into its active form depends on substrate availability, we assume saturating kinetics in this term (Eq. (A.22)). The functional form of this term is increasing for low inactive caspase-3 concentrations and saturating for large concentrations (*C_3_c__* ≫ *K*_*c*1_). Furthermore, we took into consideration the activation of caspase-3 due to the positive feedback from its active form [71], which is represented by a Hill function that gradually increases with active caspase-3 concentration (Eq. (A.23)). Here, the parameter *k*_28_ represents the strength of this positive feedback, while the Hill coefficient *n* defines the steepness of the feedback. Note that this feedback kicks in when the active caspase-3 concentration exceeds the threshold set by the Half-maximal coefficient *K*_*c*2_, which accounts for regulatory proteins that prevent caspase-3 activity (IAPs family). We also included a saturating function (Eq. (A.22)) to modulate the feedback term, accounting for substrate depletion. Furthermore, we considered the ubiquitin-protein ligase activity of XIAP that promotes the degradation of active caspase-3 [42]; therefore, it is described by Michaelis-Menten kinetics (Eq. (A.24)) similar to the degradation of p53 by Mdm2 in Eqs. (A.10) and (A.11). In this term, *k*_30_ represents the degradation rate of active caspase-3 by XIAP, while *K*_*c*3_ could be linked to the second mitochondria-derived activator of caspases (Smac) strength (not explicitly included in this description) that inhibits XIAP [46]. Finally, the parameter *k*_29_ in Eqs. (A.20) and (A.21) accounts for the basal degradation rate of active and inactive caspase-3.

The last equation of our model describes the rate of change of the concentrations of chemotherapy Doxorubicin, *D*, with respect to time,

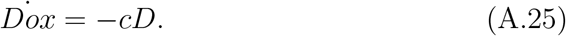

This equation considers the process of Dox clearance with a half-life between 20 to 48 hours [47]. Here, we assumed that Dox is undergoing an exponential decay with a clearance rate of *c*, and *D* is the dose level of Dox, which is considered a model input in our system.

### Appendix A.2. Model Parameters and Initial Conditions

In our model, some parameter values are based on experimental results and published models, while others are estimated. See Table. A.1 and A.2 for the model variables, initial conditions, and parameters value. In the following, we discuss a few key aspects of estimating some of our model parameters.

- p53 induces the pro-apoptotic proteins either transcriptionally in the nucleus or directly by regulating the anti-apoptotic proteins (Bcl-2 and Bcl-x) in the mitochondrial outer membrane [69]. However, Bax activation by nuclear p53 is considered to be more robust than that by cytoplasmic p53 as cells with mutated p53 (without transcriptional ability) fail to induce apoptosis. Therefore, we assumed that the Bax activation rate by cytoplasmic p53 is 12-fold less than that by nuclear p53, *k*_20_/*k*_21_ = 12.
- Considering that Mdm2 is preferentially localised in the cytoplasm [34], we assumed that the Mdm2 nuclear export rate is almost twice as much as the Mdm2 nuclear import rate. Specifically *k*_14_/*k*_13_ = 2.
- To obtain an initial rough estimate of the XIAP production rate, we first assumed that under no apoptotic conditions, the cellular concentration of XIAP depends only on its basal production and degradation rates *k*_23_ and *k*_25_, respectively. Therefore, the rate of change of XIAP concentration with respect to time can be written as: *X_c_* = *k*_23_ – (*k*_25_)(*X_c_*), then at steady state *k*_23_ = (*k*_25_)(*X_c_*). According to experimental data, the XIAP degradation rate is estimated to be 0.0116 /min, while the average basal XIAP concentration in MCF-7/C3 cells is around 0.057 mM [19]. Hence, we estimated the XIAP basal production rate to be *k*_23_ = 0.00066 mM/min.
- No experimental measurement is available for the induction rate of XIAP by Mdm2 in response to chemotherapy. Therefore, as a rough estimation, we assumed this rate to be consistent with the expected increase in the XIAP concentration following chemotherapy with respect to time as in Fig. 5a in [5].
- For caspase-3 production rate estimation, we assumed that under no apoptotic conditions, the inactive caspase-3 concentration depends only on its basal production and degradation rates *k*_26_ and *k*_29_, respectively. Then, we choose the degradation rate of inactive caspase-3 (*k*_29_=0.0039 /min) and the average inactive caspase-3 cellular concentration (0.72 mM) from the same cell type (MCF-7/C3) that we used previously to determine the rate of XIAP production [19]. Having that, we can calculate the production rate of inactive caspase-3 by assuming the inactive caspase-3 concentration at the steady state level. Thus, *k*_26_ = *k*_29_ (*C*_3_c__) = 0.0028 mM/min.
- Under apoptotic conditions, active caspase-3 is targeted by XIAP for ubiquitination, leading to caspase-3 degradation. As we do not have enough experimental data to confidently estimate this degradation rate, we arbitrarily estimated this rate (*k*_30_) to be 0.01 per min. But then we used a bifurcation analysis to study the system’s dynamical behaviour under various degradation rates of active caspase-3 by XIAP (Fig. 11a).
- In response to apoptotic stimulus, the initiator caspase-9 gets activated to process caspase-3, which in turn proteolytically cleaves and activates all caspase forms, forming a robust positive feedback loop [72]. Accordingly, we assumed that the caspase-3 activation rate by its active form is stronger than its activation by Bax by around 2-fold.

**Table A.1:** Model variables and initial conditions.

| Variable | Description | Initial Value | Unit |
| --- | --- | --- | --- |
| $ATM_n^*$ | Conc. of active nuclear ATM | 0.00 | mM |
| $p53_c$ | Conc. of cytoplasmic p53 | 0.03 | mM |
| $p53_n$ | Conc. of nuclear p53 | 0.00 | mM |
| $Mdm2_c$ | Conc. of cytoplasmic Mdm2 | 0.50 | mM |
| $Mdm2_n$ | Conc. of nuclear Mdm2 | 0.21 | mM |
| $Wip1_n$ | Conc. of nuclear Wip1 | 0.10 | mM |
| $Bax_c^*$ | Conc. of active cytoplasmic Bax | 0.00 | mM |
| $XIAP_c$ | Conc. of cytoplasmic XIAP | 0.05 | mM |
| $Casp-3_c$ | Conc. of cytoplasmic caspase-3 | 0.71 | mM |
| $Casp-3_c^*$ | Conc. of active cytoplasmic caspase-3 | 0.00 | mM |

**Table A.2:** Parameter values of the model. T.C. denotes Threshold Constant, while M.C. represents Michaelis Constant.

| Parameter | Description | Value | Unit | Reference |
| --- | --- | --- | --- | --- |
| $k_1$ | Doxorubicin-dependent $ATM_n$ activation rate | 0.001 | mM/min | Estimated |
| $K_d$ | T.C. of Doxorubicin-dependent $ATM_n$ activation | 1 | mM | [63] |
| $A_t$ | The total concentration of all $ATM_n$ forms | 1 | mM | [63] |
| $K_{at}$ | M.C. of $ATM_n$ activation | 0.2 | mM | [63] |
| $k_2$ | $ATM_n^*$ autoactivation rate | 0.095 | /min | [63] |
| $k_3$ | $Wip1_n$-dependent $ATM_n^*$ dephosphorylation rate | 0.4 | /min | [63] |
| $K_{a0}$ | M.C. of $Wip1_n$-dependent $ATM_n^*$ dephosphorylation | 0.5 | mM | [61] |
| $k_4$ | $p53_c$ basal production rate | 0.2 | mM/min | [59, 60, 61] |
| $k_5$ | $ATM_n^*$-dependent $p53_c$ phosphorylation rate | 0.2 | /min | [61] |
| $K_{p0}$ | M.C. of $ATM_n^*$-dependent $p53_c$ phosphorylation | 1 | mM | [59] |
| $k_6$ | $p53_n$ self activation rate | 0.006 | /min | Estimated |
| $K_{p1}$ | T.C. of $p53_n$ transcription activation | 0.25 | mM | [63] |
| $h$ | Hill coefficient of $p53_n$ transcription activation | 4 | - | [59, 73, 74] |
| $k_7$ | $Wip1_n$-dependent $p53_n$ dephosphorylation rate | 0.2 | /min | [59, 63] |
| $K_{p2}$ | M.C. of $Wip1_n$-dependent $p53_n$ dephosphorylation | 0.05 | mM | [63] |
| $k_8$ | $Mdm2_c$-dependent $p53_c$ degradation rate | 0.75 | /min | [61] |
| $K_{p3}$ | M.C. of $Mdm2_c$-dependent $p53_c$ degradation | 0.03 | mM | [59, 63] |
| $k_9$ | $p53_c$ basal degradation rate | 0.02 | /min | [59, 60] |
| $k_{10}$ | $Mdm2_n$-dependent $p53_n$ degradation rate | 0.02 | /min | [63] |
| $K_{p4}$ | M.C. of $Mdm2_n$-dependent $p53_n$ degradation | 0.3 | mM | [59, 63] |
| $k_{11}$ | $Mdm2_c$ basal production rate | 0.002 | mM/min | [61] |
| $k_{12}$ | $p53_n$-dependent $Mdm2_c$ production rate | 0.024 | mM/min | [59, 60, 61] |
| $k_{13}$ | $Mdm2_c$ nuclear import rate | 0.031 | /min | Estimated |
| $k_{14}$ | $Mdm2_n$ nuclear export rate | 0.07 | /min | Estimated |
| $k_{15}$ | $Mdm2_c$ basal degradation rate | 0.0028 | /min | [59, 60] |
| $k_{16}$ | $ATM_n^*$-dependent $Mdm2_n$ degradation rate | 0.0346 | /min | [63] |
| $K_m$ | M.C. of $ATM_n^*$-dependent $Mdm2_n$ degradation | 0.05 | mM | [63] |
| $k_{17}$ | $Wip1_n$ basal production rate | 0.002 | mM/min | [63] |
| $k_{18}$ | $p53_n$-dependent $Wip1_n$ production rate | 0.09 | mM/min | [61] |
| $k_{19}$ | $Wip1_n$ basal degradation rate | 0.02 | /min | [63] |
| $k_{20}$ | $p53_n$-dependent $Bax_c^*$ production rate | 0.02 | mM/min | [75] |
| $k_{21}$ | $p53_c$-dependent $Bax_c$ activation rate | 0.00167 | mM/min | Estimated |
| $K_{p5}$ | T.C. of $p53_c$-dependent $Bax_c$ activation | 0.7 | mM | Estimated |
| $h_0$ | Hill coefficient of $p53_c$-dependent $Bax_c$ activation | 3 | - | Estimated |
| $k_{22}$ | $Bax_c^*$ basal degradation rate | 0.0016 | /min | [73] |
| $k_{23}$ | $XIAP_c$ basal production rate | 0.00066 | mM/min | Estimated from [19] |
| $k_{24}$ | $Mdm2_c$-dependent $XIAP_c$ induction rate | 0.4 | /min | Estimated from [5] |
| $K_b$ | T.C. of $Bax_c^*$-dependent MOMP | 3 | mM | Estimated |
| $m$ | Hill coefficient of $Bax_c^*$-dependent MOMP | 4 | - | [75] |
| $k_{25}$ | $XIAP_c$ basal degradation rate | 0.0116 | /min | [19] |
| $k_{26}$ | $Casp-3_c$ basal production rate | 0.0028 | mM/min | Estimated from [19] |
| $k_{27}$ | $Bax_c^*$-dependent $Casp-3_c$ activation rate | 0.04 | mM/min | Estimated |
| $K_{c1}$ | M.C. of $Casp-3_c$ activation | 0.1 | mM | Estimated |
| $k_{28}$ | $Casp-3_c^*$ self activation rate | 0.07 | mM/min | Estimated |
| $K_{c2}$ | T.C. of $Casp-3_c^*$ self activation | 0.2 | mM | Estimated |
| $n$ | Hill coefficient of $Casp-3_c^*$ self activation | 4 | - | [75] |
| $k_{29}$ | $Casp-3_c$ basal degradation rate | 0.0039 | /min | [19] |
| $k_{30}$ | $XIAP_c$-dependent $Casp-3_c^*$ degradation rate | 0.01 | /min | Estimated |
| $K_{c3}$ | M.C. of $XIAP_c$-dependent $Casp-3_c^*$ degradation | 0.3 | mM | Estimated |
| $c$ | Doxorubicin clearance rate | 0.00034 | /min | Estimated from [47] |

## Data Availability

The code can be found on the Gitlab webpage https://gitlab.bham.ac.uk/spillf-systems-mechanobiology-health-disease/p53-and-xiap-dynamics.

## References

[1] S.-T. Pan, Z.-L. Li, Z.-X. He, J.-X. Qiu, and S.-F. Zhou, “Molecular mechanisms for tumour resistance to chemotherapy,” Clinical and Experimental Pharmacology and Physiology, vol. 43, no. 8, pp. 723–737, 2016.

[2] Y. Luqmani, “Mechanisms of drug resistance in cancer chemotherapy,” Medical principles and practice, vol. 14, no. Suppl. 1, pp. 35–48, 2005.

[3] R. Mirzayans, B. Andrais, P. Kumar, and D. Murray, “Significance of wild-type p53 signaling in suppressing apoptosis in response to chemical genotoxic agents: Impact on chemotherapy outcome,” International journal of molecular sciences, vol. 18, no. 5, p. 928, 2017.

[4] M. Wu, H. Ye, Z. Tang, C. Shao, G. Lu, B. Chen, Y. Yang, G. Wang, and H. Hao, “p53 dynamics orchestrates with binding affinity to target genes for cell fate decision,” Cell death & disease, vol. 8, no. 10, pp. e3130–e3130, 2017.

[5] A. L. Paek, J. C. Liu, A. Loewer, W. C. Forrester, and G. Lahav, “Cell-to-cell variation in p53 dynamics leads to fractional killing,” Cell, vol. 165, no. 3, pp. 631–642, 2016.

[6] X. Chen, J. Chen, S. Gan, H. Guan, Y. Zhou, Q. Ouyang, and J. Shi, “Dna damage strength modulates a bimodal switch of p53 dynamics for cell-fate control,” BMC biology, vol. 11, no. 1, pp. 1–11, 2013.

[7] S. Banin, L. Moyal, S.-Y. Shieh, Y. Taya, C. Anderson, L. Chessa, N. Smorodinsky, C. Prives, Y. Reiss, Y. Shiloh et al., “Enhanced phosphorylation of p53 by atm in response to dna damage,” Science, vol. 281, no. 5383, pp. 1674–1677, 1998.

[8] S.-Y. Shieh, M. Ikeda, Y. Taya, and C. Prives, “Dna damage-induced phosphorylation of p53 alleviates inhibition by mdm2,” Cell, vol. 91, no. 3, pp. 325–334, 1997.

[9] W. S. El-Deiry, “Regulation ofp53downstream genes,” in Seminars in cancer biology, vol. 8, no. 5. Elsevier, 1998, pp. 345–357.

[10] J. G. Jackson and O. M. Pereira-Smith, “p53 is preferentially recruited to the promoters of growth arrest genes p21 and gadd45 during replicative senescence of normal human fibroblasts,” Cancer research, vol. 66, no. 17, pp. 8356–8360, 2006.

[11] S. T. Szak, D. Mays, and J. A. Pietenpol, “Kinetics of p53 binding to promoter sites in vivo,” Molecular and cellular biology, vol. 21, no. 10, pp. 3375–3386, 2001.

[12] M. Kracikova, G. Akiri, A. George, R. Sachidanandam, and S. Aaronson, “A threshold mechanism mediates p53 cell fate decision between growth arrest and apoptosis,” Cell Death & Differentiation, vol. 20, no. 4, pp. 576–588, 2013.

[13] E. Batchelor, A. Loewer, C. Mock, and G. Lahav, “Stimulus-dependent dynamics of p53 in single cells,” Molecular systems biology, vol. 7, no. 1, p. 488, 2011.

[14] J. E. Purvis, K. W. Karhohs, C. Mock, E. Batchelor, A. Loewer, and G. Lahav, “p53 dynamics control cell fate,” Science, vol. 336, no. 6087, pp. 1440–1444, 2012.

[15] M. Holcik, C. Yeh, R. G. Korneluk, and T. Chow, “Translational upregulation of x-linked inhibitor of apoptosis (xiap) increases resistance to radiation induced cell death,” Oncogene, vol. 19, no. 36, pp. 4174–4177, 2000.

[16] A. Schimmer, S. Dalili, R. Batey, and S. Riedl, “Targeting xiap for the treatment of malignancy,” Cell Death & Differentiation, vol. 13, no. 2, pp. 179–188, 2006.

[17] X.-H. Yang, Z.-E. Feng, M. Yan, S. Hanada, H. Zuo, C.-Z. Yang, Z.-G. Han, W. Guo, W.-T. Chen, and P. Zhang, “Xiap is a predictor of cisplatin-based chemotherapy response and prognosis for patients with advanced head and neck cancer,” PloS one, vol. 7, no. 3, p. e31601, 2012.

[18] J. Chipuk, L. Bouchier-Hayes, and D. Green, “Mitochondrial outer membrane permeabilization during apoptosis: the innocent bystander scenario,” Cell Death & Differentiation, vol. 13, no. 8, pp. 1396–1402, 2006.

[19] J. Schmid, H. Dussmann, G. J. Boukes, L. Flanagan, A. U. Lindner, C. L. O’Connor, M. Rehm, J. H. Prehn, and H. J. Huber, “Systems analysis of cancer cell heterogeneity in caspase-dependent apoptosis subsequent to mitochondrial outer membrane permeabilization,” Journal of Biological chemistry, vol. 287, no. 49, pp. 41546–41559, 2012.

[20] Z. Sun, “The general information of the tumor suppressor gene p53 and the protein p53,” Journal of Cancer Prevention & Current Research,vol. 3, no. 1, pp. 1–13, 2015.

[21] S. Nag, J. Qin, K. S. Srivenugopal, M. Wang, and R. Zhang, “The mdm2-p53 pathway revisited,” Journal of biomedical research, vol. 27, no. 4, p. 254, 2013.

[22] Y. Haupt, R. Maya, A. Kazaz, and M. Oren, “Mdm2 promotes the rapid degradation of p53,” Nature, vol. 387, no. 6630, pp. 296–299, 1997.

[23] Y. Barak, T. Juven, R. Haffner, and M. Oren, “mdm2 expression is induced by wild type p53 activity.” The EMBO journal, vol. 12, no. 2, pp. 461–468, 1993.

[24] T. Tanaka, H. D. Halicka, F. Traganos, K. Seiter, and Z. Darzynkiewicz, “Induction of atm activation, histone h2ax phosphorylation and apoptosis by etoposide: relation to cell cycle phase,” Cell cycle, vol. 6, no. 3, pp. 371–376, 2007.

[25] A. Kurose, T. Tanaka, X. Huang, H. D. Halicka, F. Traganos, W. Dai, and Z. Darzynkiewicz, “Assessment of atm phosphorylation on ser-1981 induced by dna topoisomerase i and ii inhibitors in relation to ser-139- histone h2ax phosphorylation, cell cycle phase, and apoptosis,” Cytometry Part A, vol. 68, no. 1, pp. 1–9, 2005.

[26] C. J. Bakkenist and M. B. Kastan, “Dna damage activates atm through intermolecular autophosphorylation and dimer dissociation,” Nature, vol. 421, no. 6922, pp. 499–506, 2003.

[27] E. Meulmeester, Y. Pereg, Y. Shiloh, and A. G. Jochemsen, “Atm-mediated phosphorylations inhibit mdmx/mdm2 stabilization by hausp in favor of p53 activation,” Cell Cycle, vol. 4, no. 9, pp. 1166–1170, 2005.

[28] D. B. Young, J. Jonnalagadda, M. Gatei, D. A. Jans, S. Meyn, and K. K. Khanna, “Identification of domains of ataxia-telangiectasia mutated required for nuclear localization and chromatin association,” Journal of Biological Chemistry, vol. 280, no. 30, pp. 27587–27594, 2005.

[29] S.-H. Liang and M. F. Clarke, “Regulation of p53 localization,” European journal of biochemistry, vol. 268, no. 10, pp. 2779–2783, 2001.

[30] M. Zerfaoui, T. M. Dokunmu, E. A. Toraih, B. M. Rezk, Z. Y. Abd Elmageed, and E. Kandil, “New insights into the link between melanoma and thyroid cancer: Role of nucleocytoplasmic trafficking,” Cells, vol. 10, no. 2, p. 367, 2021.

[31] G. Gaglia, Y. Guan, J. V. Shah, and G. Lahav, “Activation and control of p53 tetramerization in individual living cells,” Proceedings of the National Academy of Sciences, vol. 110, no. 38, pp. 15497–15501, 2013.

[32] C. Amantini, P. Ballarini, S. Caprodossi, M. Nabissi, M. B. Morelli, R. Lucciarini, M. A. Cardarelli, G. Mammana, and G. Santoni, “Triggering of transient receptor potential vanilloid type 1 (trpv1) by capsaicin induces fas/cd95-mediated apoptosis of urothelial cancer cells in an atm-dependent manner,” Carcinogenesis, vol. 30, no. 8, pp. 1320–1329, 2009.

[33] K. Sakaguchi, H. Sakamoto, M. S. Lewis, C. W. Anderson, J. W. Erickson, E. Appella, and D. Xie, “Phosphorylation of serine 392 stabilizes the tetramer formation of tumor suppressor protein p53,” Biochemistry, vol. 36, no. 33, pp. 10117–10124, 1997.

[34] N. Marchenko, W. Hanel, D. Li, K. Becker, N. Reich, and U. Moll, “Stress-mediated nuclear stabilization of p53 is regulated by ubiquitination and importin-α3 binding,” Cell Death & Differentiation, vol. 17, no. 2, pp. 255–267, 2010.

[35] L. D. Mayo, J. E. Dixon, D. L. Durden, N. K. Tonks, and D. B. Donner, “Pten protects p53 from mdm2 and sensitizes cancer cells to chemotherapy,” Journal of Biological Chemistry, vol. 277, no. 7, pp. 5484–5489, 2002.

[36] N. Xu, Y. Lao, Y. Zhang, and D. A. Gillespie, “Akt: a double-edged sword in cell proliferation and genome stability,” Journal of oncology, vol. 2012, 2012.

[37] M. Fiscella, H. Zhang, S. Fan, K. Sakaguchi, S. Shen, W. E. Mercer, G. F. V. Woude, P. M. O’Connor, and E. Appella, “Wip1, a novel human protein phosphatase that is induced in response to ionizing radiation in a p53-dependent manner,” Proceedings of the National Academy of Sciences, vol. 94, no. 12, pp. 6048–6053, 1997.

[38] L. D. Mayo, Y. R. Seo, M. W. Jackson, M. L. Smith, J. R. Guzman, C. K. Korgaonkar, and D. B. Donner, “Phosphorylation of human p53 at serine 46 determines promoter selection and whether apoptosis is attenuated or amplified,” Journal of Biological Chemistry, vol. 280, no. 28, pp. 25953–25959, 2005.

[39] M. Kist and D. Vucic, “Cell death pathways: intricate connections and disease implications,” The EMBO Journal, vol. 40, no. 5, p. e106700, 2021.

[40] B. Favaloro, N. Allocati, V. Graziano, C. Di Ilio, and V. De Laurenzi, “Role of apoptosis in disease,” Aging (Albany NY), vol. 4, no. 5, p. 330, 2012.

[41] B. Dumétier, A. Zadoroznyj, and L. Dubrez, “Iap-mediated protein ubiquitination in regulating cell signaling,” Cells, vol. 9, no. 5, p. 1118, 2020.

[42] Y. Suzuki, Y. Nakabayashi, and R. Takahashi, “Ubiquitin-protein ligase activity of x-linked inhibitor of apoptosis protein promotes proteasomal degradation of caspase-3 and enhances its anti-apoptotic effect in fas-induced cell death,” Proceedings of the National Academy of Sciences, vol. 98, no. 15, pp. 8662–8667, 2001.

[43] S. Lewis and M. Holcik, “Ires in distress: translational regulation of the inhibitor of apoptosis proteins xiap and hiap2 during cell stress,” Cell death and differentiation, vol. 12, no. 6, pp. 547–553, 2005.

[44] A.-C. Godet, F. David, F. Hantelys, F. Tatin, E. Lacazette, B. Garmy-Susini, and A.-C. Prats, “Ires trans-acting factors, key actors of the stress response,” International Journal of Molecular Sciences, vol. 20, no. 4, p. 924, 2019.

[45] L. Gu, N. Zhu, H. Zhang, D. L. Durden, Y. Feng, and M. Zhou, “Regulation of xiap translation and induction by mdm2 following irradiation,” Cancer cell, vol. 15, no. 5, pp. 363–375, 2009.

[46] C. Du, M. Fang, Y. Li, L. Li, and X. Wang, “Smac, a mitochondrial protein that promotes cytochrome c-dependent caspase activation by eliminating iap inhibition,” Cell, vol. 102, no. 1, pp. 33–42, 2000.

[47] Y. L. Franco, T. R. Vaidya, and S. Ait-Oudhia, “Anticancer and cardio-protective effects of liposomal doxorubicin in the treatment of breast cancer,” Breast Cancer: Targets and Therapy, vol. 10, p. 131, 2018.

[48] A. Gabizon, H. Shmeeda, and Y. Barenholz, “Pharmacokinetics of pegylated liposomal doxorubicin,” Clinical pharmacokinetics, vol. 42, no. 5, pp. 419–436, 2003.

[49] A. Lucas, D. Lam, and P. Cabrales, “Doxorubicin-loaded red blood cells reduced cardiac toxicity and preserved anticancer activity,” Drug delivery, vol. 26, no. 1, pp. 433–442, 2019.

[50] H. Hou, D. Sun, and X. Zhang, “The role of mdm2 amplification and overexpression in therapeutic resistance of malignant tumors,” Cancer cell international, vol. 19, no. 1, pp. 1–8, 2019.

[51] M. Konopleva, G. Martinelli, N. Daver, C. Papayannidis, A. Wei, B. Higgins, M. Ott, J. Mascarenhas, and M. Andreeff, “Mdm2 inhibition: an important step forward in cancer therapy,” Leukemia, vol. 34, no. 11, pp. 2858–2874, 2020.

[52] X. Miles, C. Vandevoorde, A. Hunter, and J. Bolcaen, “Mdm2/x inhibitors as radiosensitizers for glioblastoma targeted therapy,” Frontiers in Oncology, p. 2688, 2021.

[53] F. Y. Feng, Y. Zhang, V. Kothari, J. R. Evans, W. C. Jackson, W. Chen, S. B. Johnson, C. Luczak, S. Wang, and D. A. Hamstra, “Mdm2 inhibition sensitizes prostate cancer cells to androgen ablation and radiotherapy in a p53-dependent manner,” Neoplasia, vol. 18, no. 4, pp. 213–222, 2016.

[54] L. T. Vassilev, B. T. Vu, B. Graves, D. Carvajal, F. Podlaski, Z. Filipovic, N. Kong, U. Kammlott, C. Lukacs, C. Klein et al., “In vivo activation of the p53 pathway by small-molecule antagonists of mdm2,” Science, vol. 303, no. 5659, pp. 844–848, 2004.

[55] M. Rehm, H. J. Huber, H. Dussmann, and J. H. Prehn, “Systems analysis of effector caspase activation and its control by x-linked inhibitor of apoptosis protein,” The EMBO journal, vol. 25, no. 18, pp. 4338–4349, 2006.

[56] J. R. Infante, E. C. Dees, H. A. Burris, L. Zawel, J. A. Sager, C. Stevenson, K. Clarke, S. Dhuria, D. Porter, S. K. Sen et al., “A phase i study of lcl161, an oral iap inhibitor, in patients with advanced cancer,” Cancer Research, vol. 70, no. 8 Supplement, pp. 2775–2775, 2010.

[57] L. Gu, H. Zhang, T. Liu, S. Zhou, Y. Du, J. Xiong, S. Yi, C.-K. Qu, H. Fu, and M. Zhou, “Discovery of dual inhibitors of mdm2 and xiap for cancer treatment,” Cancer cell, vol. 30, no. 4, pp. 623–636, 2016.

[58] O. Faruq, D. Zhao, M. Shrestha, A. Vecchione, E. Zacksenhaus, and H. Chang, “Targeting an mdm2/myc axis to overcome drug resistance in multiple myeloma,” Cancers, vol. 14, no. 6, p. 1592, 2022.

[59] L. Ma, J. Wagner, J. J. Rice, W. Hu, A. J. Levine, and G. A. Stolovitzky, “A plausible model for the digital response of p53 to dna damage,” Proceedings of the National Academy of Sciences, vol. 102, no. 40, pp. 14266–14271, 2005.

[60] K. B. Wee and B. D. Aguda, “Akt versus p53 in a network of oncogenes and tumor suppressor genes regulating cell survival and death,” Biophysical journal, vol. 91, no. 3, pp. 857–865, 2006.

[61] X.-P. Zhang, F. Liu, and W. Wang, “Two-phase dynamics of p53 in the dna damage response,” Proceedings of the National Academy of Sciences,vol. 108, no. 22, pp. 8990–8995, 2011.

[62] J. Elias, L. Dimitrio, J. Clairambault, and R. Natalini, “The dynamics of p53 in single cells: physiologically based ode and reaction-diffusion pde models,” Physical biology, vol. 11, no. 4, p. 045001, 2014.

[63] T. Sun, D. Mu, and J. Cui, “Mathematical model identifies effective p53 accumulation with target gene binding affinity in dna damage response for cell fate decision,” Cell Cycle, vol. 17, no. 24, pp. 2716–2730, 2018.

[64] B. Perillo, M. Di Donato, A. Pezone, E. Di Zazzo, P. Giovannelli, G. Galasso, G. Castoria, and A. Migliaccio, “Ros in cancer therapy: The bright side of the moon,” Experimental & Molecular Medicine, vol. 52, no. 2, pp. 192–203, 2020.

[65] S.-S. Li, J. Ma, and A. S. Wong, “Chemoresistance in ovarian cancer: exploiting cancer stem cell metabolism,” Journal of gynecologic oncology,vol. 29, no. 2, 2018.

[66] Y. Hirai, T. Hayashi, Y. Kubo, Y. Hoki, I. Arita, K. Tatsumi, and T. Seyama, “X-irradiation induces up-regulation of atm gene expression in wild-type lymphoblastoid cell lines, but not in their heterozygous or homozygous ataxia-telangiectasia counterparts,” Japanese journal of cancer research, vol. 92, no. 6, pp. 710–718, 2001.

[67] P. D. Jeffrey, S. Gorina, and N. P. Pavletich, “Crystal structure of the tetramerization domain of the p53 tumor suppressor at 1.7 angstroms,” Science, vol. 267, no. 5203, pp. 1498–1502, 1995.

[68] J. Chipuk and D. Green, “Dissecting p53-dependent apoptosis,” Cell Death & Differentiation, vol. 13, no. 6, pp. 994–1002, 2006.

[69] J. E. Chipuk, T. Kuwana, L. Bouchier-Hayes, N. M. Droin, D. D. Newmeyer, M. Schuler, and D. R. Green, “Direct activation of bax by p53 mediates mitochondrial membrane permeabilization and apoptosis,” Science, vol. 303, no. 5660, pp. 1010–1014, 2004.

[70] J. E. Chipuk, L. Bouchier-Hayes, T. Kuwana, D. D. Newmeyer, and D. R. Green, “Puma couples the nuclear and cytoplasmic proapoptotic function of p53,” Science, vol. 309, no. 5741, pp. 1732–1735, 2005.

[71] H. Zou, R. Yang, J. Hao, J. Wang, C. Sun, S. W. Fesik, J. C. Wu, K. J. Tomaselli, and R. C. Armstrong, “Regulation of the apaf-1/caspase-9 apoptosome by caspase-3 and xiap,” Journal of Biological Chemistry,vol. 278, no. 10, pp. 8091–8098, 2003.

[72] R. M. Reddy, W.-S. Yeow, A. Chua, D. M. Nguyen, A. Baras, M. F. Ziauddin, S. M. Shamimi-Noori, J. B. Maxhimer, D. S. Schrump, and D. M. Nguyen, “Rapid and profound potentiation of apo2l/trail-mediated cytotoxicity and apoptosis in thoracic cancer cells by the histone deacetylase inhibitor trichostatin a: the essential role of the mitochondria-mediated caspase activation cascade,” Apoptosis, vol. 12, no. 1, pp. 55–71, 2007.

[73] X. Tian, B. Huang, X.-P. Zhang, M. Lu, F. Liu, J. N. Onuchic, and W. Wang, “Modeling the response of a tumor-suppressive network to mitogenic and oncogenic signals,” Proceedings of the National Academy of Sciences, vol. 114, no. 21, pp. 5337–5342, 2017.

[74] C. Dai, H. Liu, and F. Yan, “The role of time delays in p53 gene regulatory network stimulated by growth factor,” Mathematical Biosciences and Engineering, vol. 17, no. 4, pp. 3794–3835, 2020.

[75] Z. Li, M. Ni, J. Li, Y. Zhang, Q. Ouyang, and C. Tang, “Decision making of the p53 network: Death by integration,” Journal of theoretical biology,vol. 271, no. 1, pp. 205–211, 2011.

